# A sensor complements the steric gate when DNA polymerase ε discriminates ribonucleotides

**DOI:** 10.1101/2023.07.01.547322

**Authors:** Vimal Parkash, Yashraj Kulkarni, Göran O. Bylund, Pia Osterman, Shina Caroline Lynn Kamerlin, Erik Johansson

## Abstract

The cellular imbalance between high concentrations of ribonucleotides (NTPs) and low concentrations of deoxyribonucleotides (dNTPs), is challenging for DNA polymerases when building DNA from dNTPs. It is currently believed that DNA polymerases discriminate against NTPs through a steric gate model involving a clash between a tyrosine and the 2’-hydroxyl of the ribonucleotide in the polymerase active site in B-family DNA polymerases. With the help of crystal structures of a B-family polymerase with a UTP or CTP in the active site, molecular dynamics simulations, biochemical assays and yeast genetics, we have identified a mechanism by which the finger domain of the polymerase sense NTPs in the polymerase active site. In contrast to the previously proposed polar filter, our experiments suggest that the amino acid residue in the finger domain senses ribonucleotides by steric hindrance. Furthermore, our results demonstrate that the steric gate in the palm domain and the sensor in the finger domain are both important when discriminating NTPs. Structural comparisons reveal that the sensor residue is conserved among B-family polymerases and we hypothesize that a sensor in the finger domain should be considered in all types of DNA polymerases.

## INTRODUCTION

Even though most DNA polymerases favor dNTP incorporation, the ability to discriminate ribonucleotides from deoxyribonucleotides varies among DNA polymerases (1). A conserved steric gate residue within each family of DNA polymerases has been shown to suppress the incorporation of ribonucleotides by creating a clash with the additional 2’-OH group of incoming NTPs(1–4). Family A DNA polymerases have a glutamate that functions as the steric gate, while families B and Y employ a bulky residue, such as tyrosine or phenylalanine. A third variant is found in family X polymerases that rely on a steric clash with the peptide backbone of the polymerase for sugar discrimination. This model has been confirmed in crystal structures of DNA polymerases in families A (5), X (6–8), and Y (9,10) and in structures of reverse transcriptase (3). However, until now the model for how family B DNA polymerases discriminate against ribonucleotides has relied only on biochemical and genetic experiments. These experiments have in turn been challenging because the steric gate residue, a tyrosine, is essential for efficient dNTP incorporation, and thus substitutions at this position impair DNA polymerase activity.

Studies of DNA polymerases have shown that the discrimination against ribonucleotides is not solely determined by the steric gate residue. For example, human and yeast Pol η both have a phenylalanine that acts as a steric gate residue, namely, F18 and F35, respectively (11). However, there is a dramatic difference in ribonucleotide discrimination between the two orthologues. The cause of this difference is unclear, but it might involve amino acids that influence the motions and flexibility of the active site. There are many examples of amino acid substitutions located in close proximity to the steric gate residue that influence ribonucleotide discrimination. Human Pol η has a tyrosine, Y92, that is stacked against the steric gate residue, F18(11). The Y92A substitution relaxes the ribonucleotide discrimination, presumably because the alanine gives increased flexibility of F18 that can find a position that accepts the binding of a ribonucleotide in the active site. A similar mechanism has also been proposed for other Y-family polymerases, as reviewed in (11). DNA Pol V in *Escherichia coli* has poor ribonucleotide discrimination, but the F10L substitution restricts the flexibility of the steric gate residue, Y11, and this results in increased sugar selectivity (11). Other examples where an amino acid adjacent to the steric gate residue influences ribonucleotide discrimination are found among B-family polymerases. It was first shown in the ϕ29 DNA polymerase that B-family polymerases have a conserved tyrosine (Y645 in Pol ε) that functions as a steric gate (12). The steric gate residue in Pol δ and Pol ε is essential for the incorporation of dNTPs, but substitutions introduced at the adjacent amino acid position resulted in altered discrimination against NTPs (13,14). Interestingly, the M644G substitution in Pol ε gave reduced discrimination, whereas the M644L substitution increased the discrimination, presumably by inducing either more or less flexibility, respectively, to the steric gate residue Y645 (13). An interesting report by Beese and coworkers raised the possibility that the finger domain might also be involved in recognizing misaligned ribonucleotides in the active site (5). Structures of the *Bacillus* A-family DNA polymerase I containing a ribonucleotide in the active site showed an ensemble of intermediate conformations of trapped non-cognate substrates (5). A substitution in the finger domain, F710Y, resulted in only one observed misaligned conformation of the ribonucleotide in the active site. Based on these structures with the finger domain in an ajar conformation and the biochemical characterization of the F710Y variant, it was proposed that the finger domain (the O-helix) recognizes misaligned nucleotides, including both dNTP mismatches and NTPs (5). An alternative model was later put forward by another lab which, based on studies in Family Y polymerases, proposed that a “polar filter” will discriminate ribonucleotides (15). In brief a specific residue in the finger domain was supposed to pull the bound nucleotide closer to the protein surface via hydrogen bonds with the 3’-OH group and triphosphate of the incoming nucleotide. However, when re-analyzing previously published structures we find that the geometry of B-family polymerase active sites does not convincingly support the “polar filter” model.

The *Saccharomyces cerevisiae* B-family replicative DNA polymerases, namely Pol ε, Pol δ, and Pol α, have been estimated to incorporate about 13,000 NTPs during each cell cycle of the yeast genome (13), and Pol ε seems to incorporate the majority of NTPs, roughly four times as many ribonucleotides as Pol δ. The already mentioned ^M644G^Pol ε variant has been used to address the consequences when NTPs are more frequently inserted in the genome and to determine the division of labor at the eukaryotic replication fork (13,14,16,17). This has been possible because the M644G substitution resulted in an 11-fold increase in ribonucleotide incorporation.

Here we used the ^M644G^Pol ε variant to dissect the mechanism by which a B-family DNA polymerase discriminates NTPs. A series of crystal structures were determined of ^M644G^Pol2_CORE_ (the catalytic domain of Pol ε (136 kDa)) in complex with dTTP or UTP and dCTP or CTP opposite the template dA and dG, respectively. These ribonucleotide-containing structures of ^M644G^Pol2_CORE_ showed a change in the position of the finger domain residue N828, which was in the “unlocked” orientation to accommodate a ribonucleotide. This “unlocked” position was also observed in molecular dynamics (MD) simulations of wild-type Pol2_CORE_ in complex with ATP. Biochemical and genetic studies showed that the ^N828V^Pol ε variant has an increased propensity to incorporate NTPs, suggesting that N828 functions as a sensor for discriminating against ribonucleotides. When combining the ^M644G-N828V^Pol ε variant, we observe a synergistic increase in ribonucleotide insertions, which allowed the proofreading-proficient variant of Pol ε to synthesize short stretches of RNA. The proposed mechanism might hold true for other B-family DNA polymerases because both the steric gate (Y645 in Pol ε) and sensor (N828 in Pol ε) are highly conserved residues.

## MATERIALS AND METHODS

### Expression, purification, and crystallization of Pol2__CORE__

The wild-type and exonuclease-deficient catalytic domain of *S. cerevisiae* Pol2 (Pol2_CORE_^Exo-^, residues 1-1187: D290A, E292A) and its M644G and N828V variants were expressed and purified using the protocol as described in ref. (18). Briefly, the 6× His-tagged Pol2_CORE_ was purified on a Ni^2+^-NTA column followed by overnight removal of the His-tag with PreScission protease at 6°C. The protein was passed over a Ni^2+^-NTA column a second time to remove uncleaved His-tagged protein. The protein in the flow-through was loaded onto a Mono Q column and eluted with a linear salt gradient (200–1000 mM NaAc) in 25 mM Hepes pH 7.4, 10% glycerol, and 2 mM Tris(2-carboxyethyl)phosphine hydrochloride (TCEP). The purified protein was adjusted to 25 mM Hepes pH 7.4, 800 mM NaAc, 10% glycerol, and 2 mM TCEP using a PD10 column.

For all crystallization trials, two crystallization conditions were used, either 10 mM Tris-HCl pH 8, 10 mM calcium chloride(18), and 15% PEG8000 or 50 mM MES pH6.5, 150 mM sodium acetate, 2.5% glycerol, and 8% PEG20000(19). A ternary complex with Pol2_CORE_ (and variants of Pol2_CORE_), DNA substrate (either 11ddC/16A, 11ddC/16G, or 11ddC/16T with a dA, dG, or dT opposite the incoming nucleotide, respectively), and an incoming nucleotide was formed in the presence of Ca^2+^ to inhibit the exonuclease activity when present. Crystals were flash frozen in liquid nitrogen after equilibrating the crystal in the same reservoir solution but with 15% glycerol.

A complete dataset was collected for each complex from different synchrotron beamlines (Supplementary Table S1), and the data were processed with the XDS package (20) or Mosflm (21).

### Structure determination and refinement

Phaser (22) was used to solve the structures by molecular replacement using PDB ID 4m8o (23) as the model and with one or two ternary complexes in the asymmetric unit for data processed in space groups C2 or P2, respectively. Coot (24) was used for model building, and the structures were refined using REFMAC (25) or the Phenix package (26). The refined structures had more than 95% of the residues in the most favored regions of the Ramachandran plot (Supplementary Table S1), and the model was validated using Coot (24) and MolProbity (27). PyMol (28) was used to create illustrations and to superimpose structures for comparisons.

All structures showed very good electron density in the polymerase active site region and the finger domain. Only the ^M644G^Pol2_CORE_-CTP structure (resolution 2.60 Å) lacked electron density in the finger domain (residue 790-812), distal from the polymerase active site, which might be due to a slight change in the crystal packing. However, there was no impact on the electron density in the lower part of the finger domain that is in physical contact with the incoming nucleotide.

A metal ion (Ca^2+^) was coordinated, as expected, by the catalytic aspartates (D640 and D877) in the B-site. A previously unreported Ca^2+^ ion was coordinated by the γ-phosphate of the incoming ribonucleotide and the catalytic residue D640 (not shown in Figure 1), and the Ca^2+^ ion was likely present due to the very high non-physiological concentration of Ca^2+^ ions (20mM) in the crystallization drop. This high concentration of Ca^2+^ ions was required because 20 mM ribonucleotide was added to the drop.

**Figure 1.**
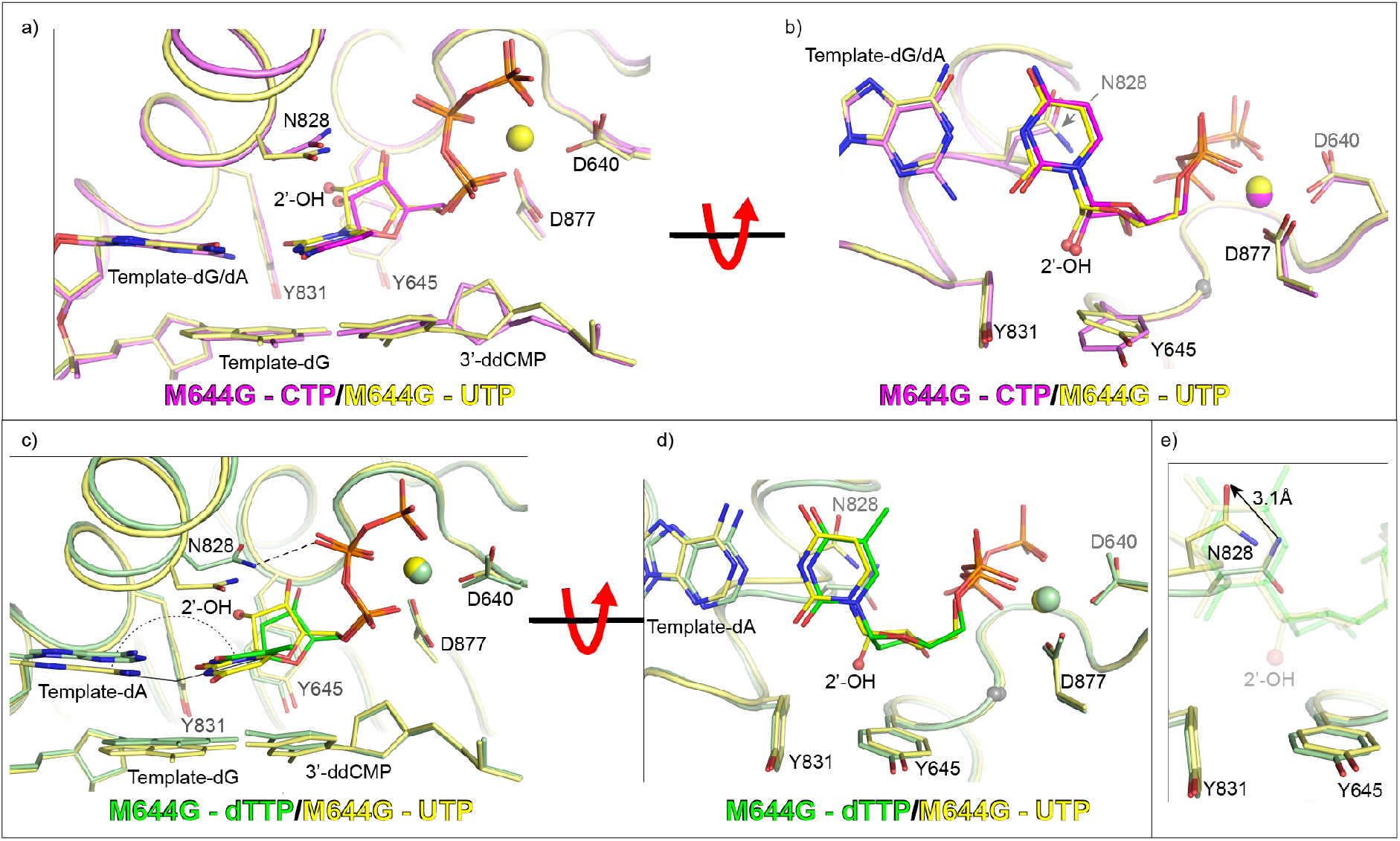
Structures of ^M644G^Pol2_CORE_ with a ribonucleotide in the active site. a) Structural superposition of ^M644G^Pol2_CORE_-CTP (in purple) and ^M644G^Pol2_CORE_-UTP (in yellow) showing how two different ribonucleotides are positioned in the active site of ^M644G^Pol2_CORE_. b) The view is rotated by 90° along the horizontal axis to show the position of the 2’-OH group relative to the steric gate, Y645. c) Comparison of ^M644G^Pol2_CORE_-UTP (in yellow) and ^M644G^Pol2_CORE_-dTTP (green) showing that the ribonucleotide takes a slightly different position compared to the deoxyribonucleotide. The nonplanar alignment with the templating base (dA) and UTP and the loose hydrogen bond between N828 and the β-phosphate of dTTP are highlighted. d) The view is rotated by 90° along the horizontal axis to show the position of the 2’-OH group relative to the steric gate, Y645. e) Zoomed in view of panel d with the incoming nucleotides in transparent sticks such that the altered position of N828 (marked with an arrow) in the ^M644G^Pol2_CORE_-UTP structure can be viewed clearly.

### Structural alignments

The structures in Supplementary Figure 1 were superimposed by aligning the Cα of residues in the Finger domain (828, 831, and 833) and in the Palm domain (645), which allowed the perfect alignment of dNTPs and showed the changes in the active site due to the M644G substitution.

The structures in Figures 1, Figure 5a, Supplementary Figure 2, Supplementary Figure 4, and Supplementary Figure 5 were superimposed by alignment of the Palm domain residues 877, 640, 556, and 645, and one residue from the Finger domain, 836.

The structures in Figure 5b were superimposed by alignment of the Palm domain residues 640, 644, 645, and 556, and the Finger domain residue 836.

### MD simulations

MD simulations were performed on wild-type Pol2_CORE_ (PDB ID: 4m8o (23)) and ^M644G^Pol2_CORE_ (PDB ID: 6fwk) in complex with dATP or ATP. Each structure, in addition to the polymerase in complex with the nucleotide, also consisted of one Mg^2+^ in the active site and one Zn^2+^ ion at the base of the P-domain. The Mg^2+^ and Zn^2+^ ions were replaced by dummy models using the parameters described in ref. (29). This allowed us to capture the electrostatic and structural properties of the ions without requiring artificial bonds or restraints. In the case of the Mg^2+^ ion, we used an octahedral model taking parameters directly from ref. (29). In the case of the Zn^2+^ ion, we constructed a tetrahedral model using the non-bonded parameters from ref. (29) (Supplementary Table 5). This model was used to capture the geometric properties of the tetrahedrally coordinated Zn^2+^ ion present in the crystal structure (Supplementary Figure 7 and Supplementary Table 6). Coordinates of the bound incoming nucleotide in the structure were used to generate both dATP-bound and ATP-bound models.

All MD simulations were performed using the AMBER18 simulation package (30) and followed the protocol described in ref. (19). The ff14SB force field (31,32) was used to describe the protein, and the Parmbsc1 force field (33) as implemented in AMBER18 was used to describe the DNA. The initial minimization, equilibration, and production runs were carried out using the PMEMD module (34), and LEaP was used to generate the topology and initial coordinates. An octahedral solvent box comprised of TIP3P water molecules (35) was used to solvate the protein-DNA complex, with the box extending at least 10 Å from the solute in either direction. Aspartate, glutamate, lysine, and arginine residues were kept in their ionized states, while histidine residues were singly protonated at the Nε atom. The total charge of the system was neutralized by adding the requisite number of Na^+^ counter ions. Additionally, Na^+^ and Cl^−^ ions were added to the system to maintain an ionic strength of 0.15 M.

Upon solvation the system was subjected to an energy minimization procedure, which consisted of 2000 steps of steepest descent followed by 3000 steps of conjugate gradient minimization using 5 kcal mol^−1^ Å^−2^ harmonic positional restraints on all heavy atoms of the solute. The following equilibration protocol was then used. 1; NVT simulations of 1.5 ns were used to increase the temperature of the system in three 500 ps steps from 5 K to 100 K in the first step, from 100 K to 200 K in the second step, and from 200 K to 300 K in the third step. 2; A 1 ns NPT equilibration at a constant isotropic pressure of 1 atm in five steps of 200 ps each was used to decrease the harmonic positional restraints on the solute heavy atoms progressively from 5 kcal mol^−1^ Å^−2^ to 1 kcal mol^−1^ Å^−2^. Finally, 3; a 500 ps NPT simulation without any restraints on the system was run. This resulted in an overall equilibration time of 3 ns. All simulations were performed using the Berendsen thermostat (36) and pressure control algorithms using a 1 ps time constant.

The end point of the equilibration step was used to initiate three independent 130 ns production simulations of both the wild-type and M644G variants (bound to either dATP or ATP). This resulted in three simulation replicas for each variant, each initiated using different starting velocities. Thus, each variant was sampled for a total simulation time of 390 ns per system, giving a total simulation time of 1.56 µs over all systems. NPT conditions were used, and a constant temperature of 300 K was maintained using the Langevin thermostat (37) with a collision frequency of 2 ps^−1^. A constant pressure of 1 atm was maintained using the Monte Carlo barostat (38), and the SHAKE algorithm (39,40) was used to constrain all bonds involving hydrogen atoms. Short-range non-bonded interactions were calculated using a 10 Å cutoff radius. Long-range electrostatic interactions were described using the particle mesh Ewald method (41,42), and all simulations were performed using a 1 fs time step with frames saved every 2.5 ps.

### Analysis

The final 100 ns of each production simulation was used for the analysis, including the measurement of interatomic distances and angles. This was done with the intention of allowing the system to equilibrate further for 30 ns before using the simulation to perform the analysis. This resulted in 300 ns (3 × 100 ns) of simulation data per system available for analysis. For all analyses of simulation data, frames were extracted at every 12.5 ps of the trajectory, leading to the extraction of a total of 24,000 frames per system.

Clustering analysis on the MD trajectories was performed using CPPTRAJ (43). The Hierarchical-Agglomerative clustering algorithm was used with a sieve value of 5 and a cutoff distance of 2.0 Å. Measurements of interatomic distances and angles were performed using VMD 1.9.1 (44).

### Expression and purification of four-subunit Pol ε

*S. cerevisiae* wild-type Pol ε, Pol ε exo^−^ (pol2^D290A, E292A^), and its variants (N828V, M644G and M644G,N828V) were over-expressed in *S. cerevisiae* as described in ref. (45). Wild-type, exonuclease proficient Pol ε, ^N828V^Pol ε, ^M644G^Pol ε, and ^M644G,N828V^Pol ε were purified as described in ref. (45). Exonuclease-deficient Pol ε, ^N828V^Pol ε, ^M644G^Pol ε, and ^M644G,N828V^Pol ε were purified with the help of a FLAG-tag on Pol2 as described in ref. (18).

### Primer extension assays

The primer extension assays were performed essentially as described in ref. (23). Briefly, a 10 μL reaction mix A (10 nM Pol ε and 10 nM primer/template, 20 mM Tris-HCl pH 7.8, 20 mM NaAc, 0.1 mg/mL bovine serum albumine, and 0.5 mM dithiothreitol) was pre-incubated on ice then mixed with 10 μL reaction mix B (16 mM MgAc2, 20 mM Tris-HCl pH 7.8, 0.1 mg/mL bovine serum albumin, and 0.5 mM dithiothreitol and with physiologically balanced dNTPs or dNTPs/NTPs at 2× the indicated concentration) and incubated for the indicated time at 30°C. The reactions were stopped by the addition of 20 μL stop solution (96% formamide, 20 mM EDTA, and 0.1% bromophenol blue). The reactions were heated to 90°C for 15 minutes before the products were separated on a denaturing polyacrylamide gel. The primer strand was labeled at the 5′ end by tetrachlorofluorescein (TET) to allow detection of products.

### *In vitro* incorporation of rNMPs into DNA

Stable incorporation of rNMPs by the replicative polymerases was analyzed using a substrate made by annealing a 50-mer ^32^P-labeled primer strand to an 80-mer template strand. The reaction mixtures contained 100 nM DNA substrate, and the reaction buffer for each polymerase was as described for primer-extension reactions (see above). Nucleotides were added at cellular concentrations (3 mM ATP, 1.7 mM UTP, 0.5 mM CTP, 0.7 mM GTP, 16 μM dATP, 30 μM dTTP, 14 μM dCTP, and 12 μM dGTP) (46) and included only four dNTPs or all eight nucleotides (all four dNTPs and all four NTPs). Reactions were initiated by adding 35.5 nM Pol ε, followed by incubation at 30°C and termination after 30 min by the addition of an equal volume of formamide with loading dye. The products were separated on a denaturing 10% polyacrylamide gel, and the positions of full-length products were identified by a brief exposure on X-ray film followed by excision from the gel and purification. Equivalent amounts of purified DNA (as determined by scintillation counting) were treated with either 0.3M KCL or 0.3M KOH for 2 h at 55°C. Following addition of an equal volume of formamide with loading dye, equivalent amounts of samples were analyzed by electrophoresis on a denaturing 10% polyacrylamide gel. Products were detected using a phosphorImager and ImageQuanT software.

### *In vivo* analysis

A linearized integration plasmid p173 (47) carrying mutations resulting in *pol2 N828V* or pol2 *M644G,N828V* was integrated into a diploid E134 yeast strain (*ade5-1 lys2::InsEA14 trp1-289 his7-2 leu2-3,112 ura3-52*). Four integrants from each transformation were isolated and then patched on YPD overnight to allow for the looping out of the URA3 marker, leaving the specific POL2 mutation on the chromosome. Patched clones were then printed on 5-FOA plates to select for clones that had lost URA3. Three 5-FOA^r^ clones from each patch were picked and streaked for single cells on YPD. The respective mutations were screened by PCR, and positive diploid clones were sequenced across the *POL2* gene to confirm that the selected mutation was correctly integrated into the heterozygous strain. At least 8 tetrads from each strain were dissected as previously described (48).

### Detection of incorporated ribonucleotides *in vivo*

The strains used to monitor ribonucleotide incorporation, namely *pol2-M644G Δrnh201::KanMX* and *pol2-N828V Δrnh201::KanMX*, were constructed by first inserting a *pol2-M644G* or *pol2-N828V* mutation into the diploid E134 strain as described (18) to create a heterozygous *POL2/pol2-M644G* strain and a *POL2/pol2-N828V* strain. Each strain was next transformed with a PCR-amplified fragment with a *KanMX* cassette that was integrated into the *RNH201* gene to generate the diploid heterozygous *POL2/pol2-M644G, RNH201/Δrnh201::KanMX* strain and the *POL2/pol2 N828V, RNH201/Δrnh201::KanMX* strain. Finally, the diploid strains were sporulated, and haploid *pol2-M644G Δrnh201::KanMX* and *pol2-N828V Δrnh201::KanMX* strains were isolated. Genomic DNA was extracted from wild-type E134, *pol2-M644G Δrnh201::KanMX*, and *pol2-N828V Δrnh201::KanMX* and was treated with alkali to hydrolyze sites where ribonucleotides were incorporated in the DNA. The samples were analyzed by southern blot followed by hybridization with a radioactive probe, essentially as described in refs. (13,17,49) except that the ^32^P-labelled single-stranded probe was complementary to either the lagging strand or the leading strand at the *FUS1* gene, approximately 1 kb from the origin ARS306 on chromosome III.

### Flow cytometry

The cell cycle distribution in an asynchronous cell culture was measured essentially as described in ref. (46) A total of 4 ml of asynchronously growing cells were harvested in logarithmic phase at A600 = 0.3 and fixed overnight in 70% ethanol at 4°C. Fixed cells were washed in water followed by incubation in 0.5 ml of 2 mg/ml RNase A, 50 mM Tris-HCl pH8.0, and 15 mM NaCl at 37°C for 15 h. RNase A was inactivated by the addition of 50 μl (20 mg/ml) Proteinase K, and the sample was incubated at 50°C for 75 min. The RNase A-treated cells were pelleted at 2500 × *g* for 3 min and resuspended in 0.5 ml of 50 mM Tris-HCl pH7.5, and 50 μl of resuspended cells were stained in 1 ml SYBR Green solution (Molecular Probes, SYBR Green diluted 1:10,000 in 50 mM Tris-HCl pH7.5) and sonicated for 10 seconds with a QSONICA Q500 sonicator equipped with a micro tip and with the amplitude set to 20% prior to analysis in a Cytomics FC500 flow cytometer (Beckman Coulter).

## RESULTS

### Two crystal structures of ^M644G^Pol2_CORE_ with a ribonucleotide in the active site

Initial attempts were made to obtain a ternary structure of the catalytic domain of wild-type Pol ε (Pol2_CORE_) with a ribonucleotide in the active site, but no protein crystals were formed. Previous biochemical experiments showed that ^M644G^Pol ε has an increased propensity to incorporate NTPs (13), suggesting that ^M644G^Pol ε might more easily accommodate a ribonucleotide in the active site. Thus, we mixed ^M644G^Pol2_CORE_ with a primer-template and a very high concentration of a ribonucleotide (see Materials and Methods). Two different ternary complex structures of ^M644G^Pol2_CORE_ with UTP or CTP (^M644G^Pol2_CORE_-UTP (PDB ID 8b79) and ^M644G^Pol2_CORE_-CTP (PDB ID 8b67)) were solved at 2.65 Å and 2.60 Å, respectively (Figure 1 and Supplementary Table 1). The two structures of ^M644G^Pol2_CORE_-UTP and ^M644G^Pol2_CORE_-CTP aligned well in the polymerase active site when the two structures were superimposed (Figures 1a-b). In both structures, the NTPs adopted the same C3’-endo conformation, and the 3’-dideoxy end of the primer terminus was at the same distance from the α-phosphate of the incoming nucleotide as found in structures with a dNTP. Thus, the NTPs were in a position admissible for chemistry to occur if a 3’-OH group were to be present at the DNA primer terminus.

### Comparison of the structures of M644GPol2_CORE_ and wild-type Pol2_CORE_ suggests that the M644G substitution introduces flexibility in the polymerase active site

To clarify the impact of M644G and NTPs on the geometry of the active site, we solved two new structures of ^M644G^Pol2_CORE_ with a deoxyribonucleotide, including ^M644G^Pol2_CORE_-dCTP (PDB ID: 8b6k) and ^M644G^Pol2_CORE_-dTTP (PDB ID: 8b76, Supplementary Table 1) in order to complement the already published structures of Pol2_CORE_-dATP (PDB ID: 4m8o (23), PDB IDs: 6h1v and 6qib (18)), Pol2_CORE_-dCTP (PDB ID: 4ptf (50)), and ^M644G^Pol2_CORE_-dATP (PDB: 6fwk (19)). First, ^M644G^Pol2_CORE_-dATP (PDB ID: 6fwk) was compared with Pol2_CORE_-dATP (PDB ID: 4m8o) and was found to have an almost identical active site geometry as that of Pol2_CORE_-dATP. Specifically, only a small shift of the catalytic residue 877 was observed, likely due to the void volume between Y645 and the catalytic residue D877 in ^M644G^Pol2_CORE_ (Supplementary Figure 1a). Aligning all available wild-type Pol2_CORE_ structures (PDB IDs: 4m8o, 4ptf, 6qib and 6h1v) and all ^M644G^Pol2_CORE_ structures with a dNTP in the active site confirmed that there were, overall, no major changes in the ^M644G^Pol2_CORE_ structures. However, a small ∼1Å shift in the backbone Cα-atom of the catalytic residue D877 towards the cavity between D877 and Y645 was consistently observed in all ^M644G^Pol2_CORE_ structures when compared to wild-type Pol2_CORE_ structures (Supplementary Figures 1b and 1c). This consistent structural shift in D877 suggests increased flexibility at the polymerase active site of ^M644G^Pol2_CORE_, something that was hypothesized when ^M644G^Pol2_CORE_ was created for use in earlier studies (13,14,16,17,47).

### The position of the finger domain residue N828 is altered in the presence of ribonucleotides

The comparison between the UTP and dTTP-bound structures of ^M644G^Pol2_CORE_ showed that the position of C2’ in the sugar of the ribonucleotide was slightly shifted upward and sideways toward the finger domain such that the 2’-OH group of the ribonucleotide was positioned between Y645 and Y831 (Figures 1c and 1d) without altering the position of the steric gate residue Y645. Consequently, the base of the incoming ribonucleotide was slightly tilted, and the nascent dA:UTP base-pair became non-planar (Figure 1c). The slight shift of the sugar pucker towards the finger domain resulted in a large shift in the position of N828, which is a residue in the finger domain (Figures 1c and 1e). Furthermore, a local distortion around N828 in the α-helix of the finger domain suggests that the new position of N828 induced a strain in the α-helix. The described structural changes were also found when comparing ^M644G^Pol2_CORE_-CTP with ^M644G^Pol2_CORE_-dCTP (Supplementary Figure 2).

### MD simulations

To investigate whether the N828 conformational shifts observed in the ^M644G^Pol2_CORE_-UTP crystal structure also occurred in wild-type Pol2_CORE_-NTP, we performed MD simulations of wild-type Pol2_CORE_ and ^M644G^Pol2_CORE_ in complex with either dATP or ATP. During these simulations, we tracked the structural dynamics of three amino acid side chains – Y645, N828, D877 – and how they were impacted by the presence of the incoming nucleotide. The crystal structures of ^M644G^Pol2_CORE_ showed that a truncation of the M644 side chain to glycine led to a spatial shift in the active site residue D877 (Supplementary Figure 1c). The distances between the Cα atom of the D877 side chain and the hydroxyl group of the Y645 side chain were measured in all the systems to check if the space created by the M644G substitution led to a shorter distance between the D877 and Y645 side chains, as observed in the crystal structures.

Distances averaged over all simulations showed that the distance between the D877 and Y645 side chains was reduced in ^M644G^Pol2_CORE_ compared to wild-type Pol2_CORE_ (Supplementary Figure 6 and Supplementary Table 2). Thus, the observed shift in the protein crystals was likely the result of increased flexibility in the active site.

To further understand the influence of Y645 on the recognition of the incoming nucleotide in the various Pol ε systems, the distance between C_δ1_ of Y645 and 2’-C of the incoming nucleotide (referred to here as the “sugar ring - Y645 distance”) was measured. The analysis showed that in both ^M644G^Pol2_CORE_ and wild-type Pol2_CORE_ the distance between the specific atoms of Y645 and the sugar ring was consistently greater by ∼0.6 Å for a ribonucleotide compared to a deoxyribonucleotide when bound to the active site (Supplementary Figure 6 and Supplementary Table 3). Thus, the clash between the 2’-OH of the ribonucleotide and the steric gate Y645 resulted in an altered position of the ribose (Supplementary Figure 2). The shift in the position of the finger domain residue N828 was also investigated. Based on the observed shift of the N828 residue in the crystal structures, the angle formed by the Cα and Cγ atoms of N828 and the O2B oxygen bound to the β-phosphate of the incoming nucleotide (referred to as the “N828 shift” below) was measured (Figures 2e-f). Here, a wider angle (closer to 160°) would indicate that the N828 side chain was in a “locked” position, which was observed in the dNTP-bound crystal structure of ^M644G^Pol2_CORE_ (as found in PDB ID 8b76, Figure 2e)). A decreased angle would represent the “unlocked” conformation of N828, as was observed in the ribonucleotide-bound crystal structures of ^M644G^Pol2_CORE_ (PDB ID 8b79, Figure 2f). The average values of this angle over all simulation replicas for wild-type Pol2_CORE_ and ^M644G^Pol2_CORE_ bound to dATP/ATP confirmed the trend observed in the crystal structures that N828 underwent a conformational “outward” shift when a dNTP was replaced by a ribonucleotide in the active site (Supplementary Figure 6 and Supplementary Table 4).

**Figure 2.**
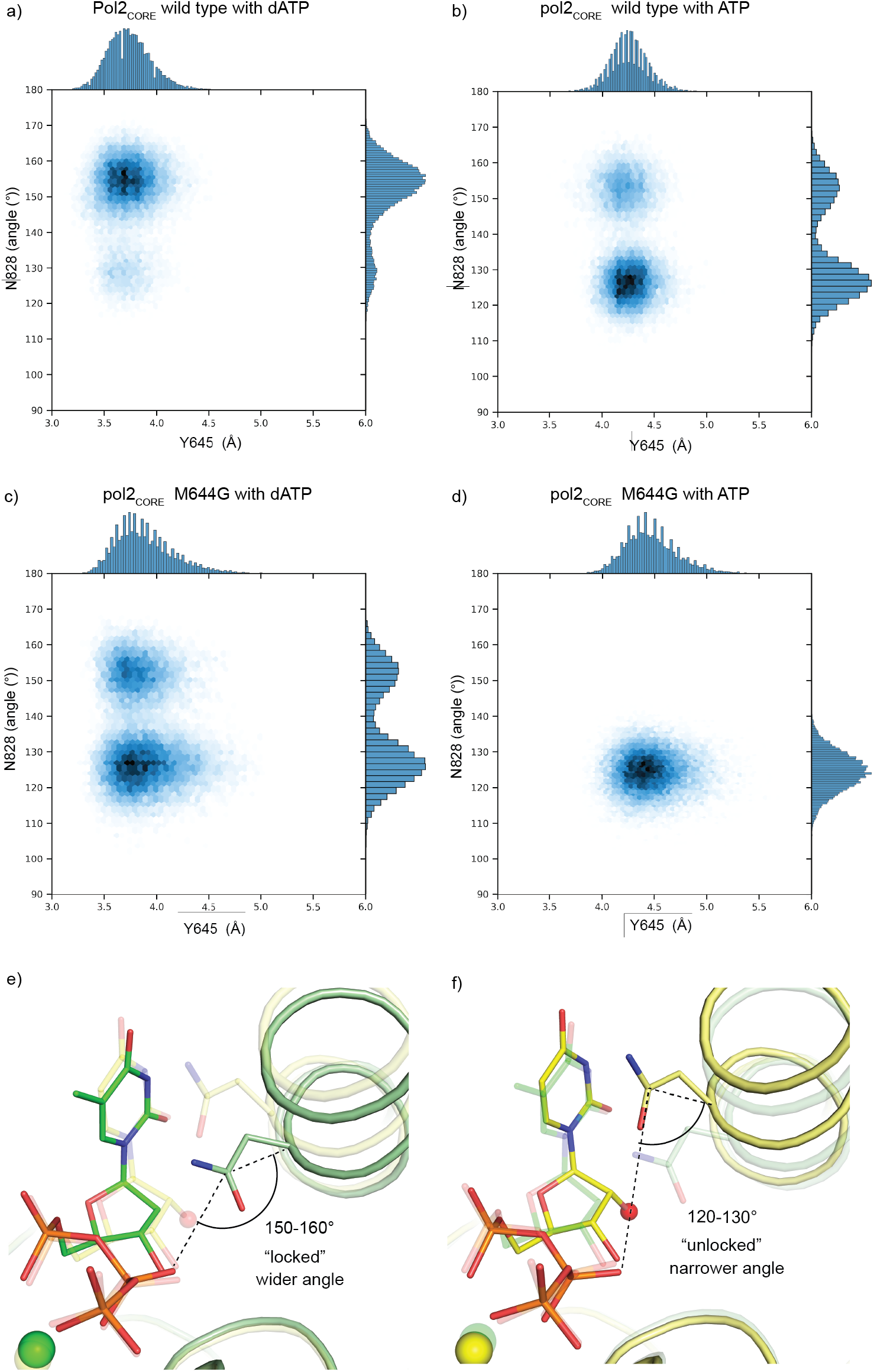
MD simulations of wild-type Pol2_CORE_ and ^M644G^Pol2_CORE_ in complex with either dATP or ATP. a) Wild-type Pol2_CORE_ with a bound dATP. b) Wild-type Pol2_CORE_ with a bound ATP. c) ^M644G^Pol2_CORE_ with a bound dATP. d) ^M644G^Pol2_CORE_ with a bound ATP. The distance between C81 of Y645 and the 2’-C of the incoming nucleotide is plotted on the horizontal axis. To monitor the position of N828, the angle formed by the Cα and Cγ atoms of N828 and the O2B oxygen of the β-phosphate in the nucleotide (see illustrations in panel e and f) is plotted on the vertical axis. e) Illustration showing the wider angle observed in the crystal structure of ^M644G^Pol2_CORE_ in complex with dTTP (green). f) Illustration showing the narrower angle observed in the crystal structure of ^M644G^Pol2_CORE_ in complex with UTP (yellow). Note that the angles in panel e) and f) correspond well with the favored position of N828 in the MD simulations shown in panels a-d.

Further analysis of the correlation between shifts in Y645 and N828 was performed by plotting a 2D scatter plot of the “sugar ring - Y645 distance” on the X-axis and the N828 angle on the Y-axis (Figures 2a-d). In wild-type Pol2_CORE_, the distance between Y645 and the 2’-C of the incoming nucleotide increased when an ATP was bound (compare panels a and b). In the presence of dATP, a position with a wider angle dominated (the locked conformation), whereas in the presence of ATP a shift was observed towards a narrower angle (the unlocked conformation). Thus, N828 was less frequently able to form a stable interaction with the β-phosphate of the incoming nucleotide if there was a ribonucleotide in the active site. A similar observation was made in ^M644G^Pol2_CORE_ (Figure 2, compare panels c and d). In fact, the MD simulations suggested that N828 in ^M644G^Pol2_CORE_ was less prone to form a loose hydrogen bond with the β-phosphate even when a dATP was bound to the active site (Figure 2, compare panels a and c). This could be explained by increased flexibility in the active site that may affect the position of the incoming nucleotide and result in the observed reduced fidelity of ^M644G^Pol ε^19^. In the presence of ATP, the narrower angle of N828 was exclusively observed, as shown in the structures in Figure 1c and 1e.

In conclusion, both the MD simulation data and the structural data confirmed that Y645 acts as a steric gate due to a clash with the 2’-OH of the ribonucleotide, which results in an increased distance between Y645 and the sugar ring. As a consequence, two preferred positions were observed for the side-chain of N828, namely a locked position stabilized by a loose hydrogen bond to the β-phosphate when a deoxyribonucleotide was bound and an unlocked position, which was not stabilized by the hydrogen bond, when a ribonucleotide was bound.

### Ribonucleotide incorporation by N828VPol ε

To test the hypothesis that steric hindrance does not allow the finger domain to adopt a completely closed conformation when a ribonucleotide is bound to the active site, we replaced N828 with a valine, which is both less bulky and unable to form a hydrogen bond with the β-phosphate of the incoming nucleotide. First, we asked whether the polymerase activity was affected in a primer extension assay with physiological concentrations of dNTPs. The activity of ^N828V^Pol ε was comparable to wild-type Pol ε and ^M644G^Pol ε (Figure 3a). When repeating the experiment with a physiological concentration of both dNTPs and NTPs, the polymerase activity of wild-type Pol ε was suppressed due to the presence of high concentrations of NTPs. The ^M644G^Pol ε variant was less suppressed by NTPs, and ^N828V^Pol ε was even less affected (Figure 3a), suggesting that ^N828V^Pol ε is more tolerant of NTPs. However, the assay did not reveal to what extent NTPs were incorporated. To investigate if ^N828V^Pol ε incorporates NTPs at physiological concentrations of dNTPs/NTPs, the primer extension was continued for a longer period of time such that all products were full-length products. The full-length products were extracted and treated with either KCl or KOH at high temperature before separating the fragmented DNA on a sequencing gel. The shorter products after treatment with KOH were the result of hydrolysis at positions where NTPs were incorporated during the primer-extension reaction. ^N828V^Pol ε, like the ^M644G^Pol ε mutant, incorporated more NTPs than Pol ε (Figure 3b). In addition, the pattern of the products suggested that ^N828V^Pol ε and ^M644G^Pol ε have different preferences for sequence context when incorporating NTPs.

**Figure 3.**
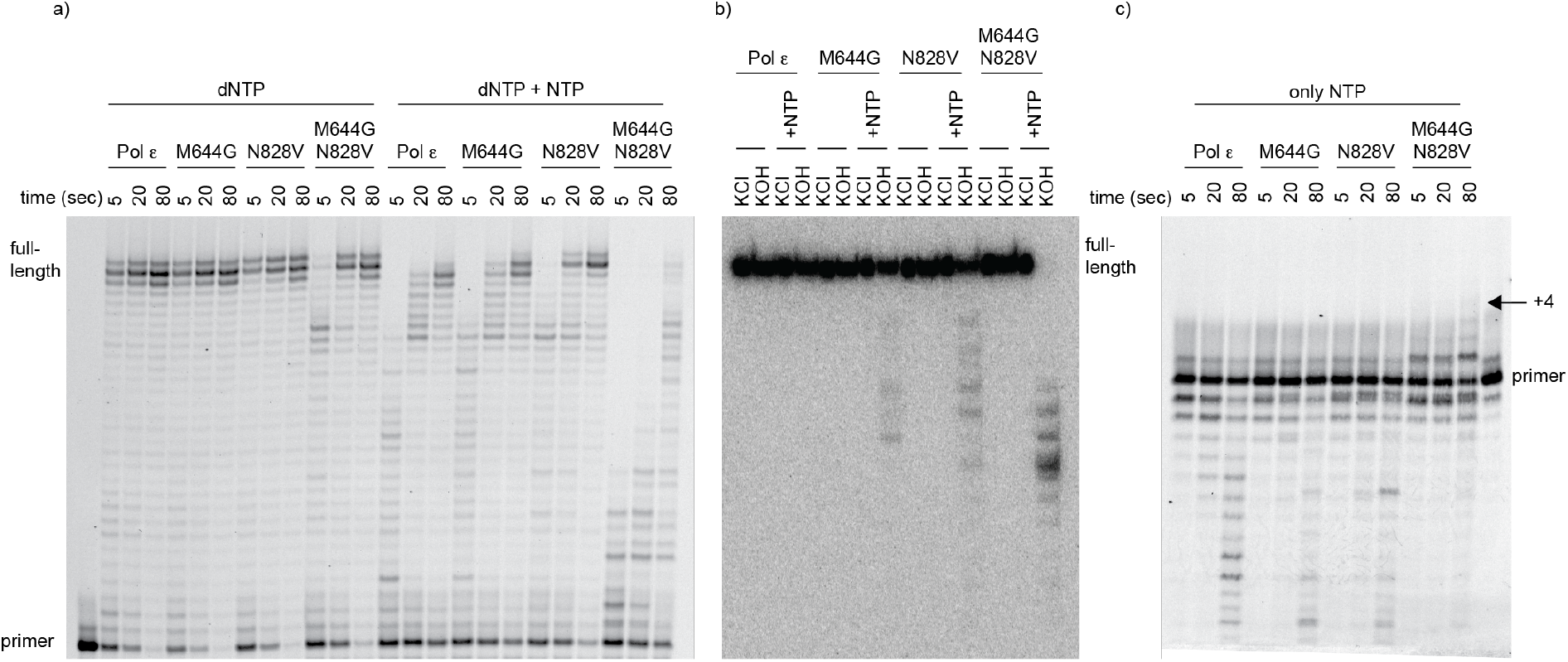
^N828V^Pol E incorporates ribonucleotides more frequently in the nascent strand compared to wild-type Pol ε. a) Primer extension assays with wild-type Pol ε, ^M644G^Pol E, ^N828V^Pol2 E, and ^M644G-N828V^Pol E were performed by mixing preformed enzyme-DNA complexes with magnesium acetate and physiological concentrations of dNTPs alone or dNTPs mixed with NTPs. Reactions were incubated for 5, 20, and 80 sec at 30°C and terminated products were separated on a 10% denaturing acrylamide gel. ^N828V^Pol E was more tolerant to the addition of physiological levels of NTPs when compared to wild-type Pol E. Only ^M644G-N828V^Pol E showed decreased DNA polymerase activity in the presence of dNTPs and reduced capacity to build long products in the presence of physiological levels of NTPs. b) Fully extended replication products (30 nucleotides) by exonuclease-deficient Pol ε, ^M644G^Pol E, ^N828V^Pol2 E, and ^M644G-N828V^Pol E were treated with either KCl or KOH at high temperature in order to induce breaks in the DNA strand where NTPs were incorporated. Shorter products were obtained when products extended by ^M644G^Pol E, ^N828V^PolE, or ^M644G-N828V^Pol E were incubated with KOH. All products built by ^M644G-N828V^Pol E included at least one ribonucleotide. c) Primer extension assays with wild-type Pol ε, ^M644G^Pol E, ^N828V^Pol2 E, and ^M644G-N828V^Pol E were performed by mixing preformed enzyme-DNA complexes with magnesium acetate and physiological concentrations of only ribonucleotides. Reactions were incubated for 5, 20, and 80 sec at 30°C, and terminated products were separated on a 10% denaturing acrylamide gel. Within 80 seconds, proofreading-proficient ^M644G-N828V^Pol E added four consecutive NTPs to the nascent strand.

To explore if the combination of M644G and N828V would increase ribonucleotide incorporation even further, the ^M644G-N828V^Pol ε variant was expressed and purified. Surprisingly, all primer-extension products that were extended by ^M644G-N828V^Pol ε contained at least one ribonucleotide despite the extended strand being only 30 nt long (Figure 3b).

However, the capacity to build DNA was reduced when only dNTPs and when both dNTPs and NTPs were added to the reaction (Figure 3a). Interestingly, with only NTPs in the reaction, proofreading-proficient ^M644G-N828V^Pol ε could insert up to 4 consecutive NTPs after 80 seconds and thus could be considered to be a RNA polymerase (Figure 3c). In contrast, proofreading-proficient wild-type Pol ε, ^M644G^Pol ε, and ^N828V^Pol ε were unable to insert consecutive NTPs and instead degraded the primer (Figure 3c).

To determine whether ^N828V^Pol ε and ^M644G-N828V^Pol ε also incorporate more ribonucleotides when replicating DNA *in vivo*, yeast strains with these mutants were constructed in a diploid E134 background. After sporulation, haploid strains were isolated as either *RNH201* or *rnh201Δ*, expressing wild-type Pol ε, ^M644G^Pol ε, ^N828V^Pol ε, or ^M644G-N828V^Pol ε. Yeast strains with an *rnh201Δ* are unable to remove ribonucleotides from the genome. The first observation was that *RNH201, pol2-M644G,N828V* haploid strains were very sick strains, so further downstream analyses were not possible due to the high probability of suppressor mutations (Supplementary Figure 3a). In contrast, the *pol2-N828V* strains were viable, with a doubling time similar to *pol2-M644G* strains (Supplementary Figure 3b and Figure 4a). Both strains had a slightly longer doubling time than wild-type strains, and flow cytometry showed that exponentially growing cultures had a slightly increased S and G2/M population (Figure 4a-b). Next, M644G and N828V were combined with *rnh201Δ* in a heterozygous diploid strain, and cells were isolated as haploids after sporulation. The *pol2-M644G rnh201Δ*, *pol2-N828V rnh201Δ*, and *POL2 rnh201Δ* strains were then harvested during the exponential growth phase. The genomic DNA was isolated and treated under alkaline conditions at high temperature to induce single-strand breaks at sites where ribonucleotides were incorporated. The fragments were separated on an alkaline agarose gel followed by hybridization of a radiolabeled probe to the nascent leading strands. ^N828V^Pol ε incorporated ribonucleotides when synthesizing the leading strand, as previously shown for ^M644G^Pol ε, resulting in shorter fragments compared to the wild-type Pol ε strain (Figure 4c). When hybridizing a radiolabeled probe complementary to the newly synthesized lagging strand, both strains showed only longer products (Figure 4c). These results supported the model that Pol ε primarily participates in leading strand synthesis.

**Figure 4.**
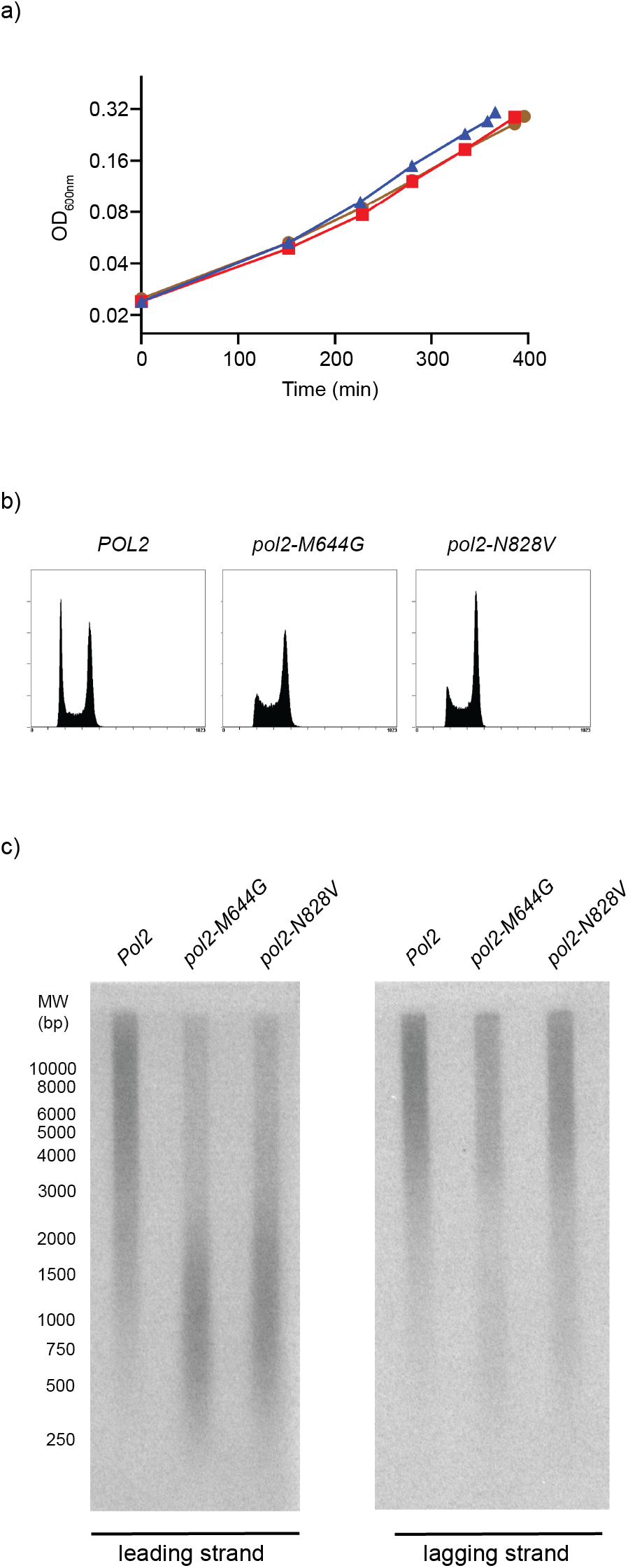
Haploid yeast expressing ^N828V^PolE have similar phenotypes as strains expressing ^M644G^Pol E and incorporate ribonucleotides during leading strand synthesis. a) Growth curves for haploid yeast strains with a *pol2-N828V* (red squares) or a *pol2-M644G* (brown circles) allele showing similar doubling times, which are not much slower than the parental wild-type *POL2* strain (blue triangles). b) Yeast strains with a *pol2-N828V* or a *pol2-M644G* allele showed a similar cell cycle distribution in asynchronous cultures, with a slight increase in S and G2/M cells and a reduction in G1 cells when compared to the parental wild-type strain. c) Haploid *rnh201Δ* strains with either a *pol2-N828V* allele (see Figure 4b) or a *pol2-M644G* allele were isolated. Genomic DNA was extracted and treated with alkali followed by Southern blotting, and a single-stranded radiolabeled probe was hybridized to either the nascent leading or lagging strand.

The reduced discrimination against NTPs may also suggest an increased mutation rate due to replication errors such as mismatches. This was previously shown to be the case for *pol2-M644G*, which showed an increased error rate for T-T mismatches (47). Thus, the spontaneous mutation rates for the wild-type and *pol2-N828V* strains were measured in the forward mutation assay using the *CAN1* locus, and this showed a 3.8-fold elevated mutation rate in the *pol2-N828V* strain (Supplementary Table 7). Next, we attempted to establish a *pol2-N828V msh2Δ* strain with a complete loss of mismatch repair, but the strain’s growth was impaired both in liquid culture and on plates. Thus, these strains were likely accumulating suppressor mutations. Instead, we combined *pol2-N828V* with *msh6Δ*, which showed only a partial loss of mismatch repair. We found a synergistic increase with the loss in mismatch repair suggesting that the bulk of errors originated during DNA replication when *pol2-N828V* synthesized the leading strand (Supplementary Table 7). Further sequence analysis of mutations in *CAN1* in independent isolates of *pol2-N828V* strains revealed an even stronger prevalence for mutations that most likely originated from T-T mismatches on the leading strand compared to the *pol2-M644G* (*47*) strain (Supplementary Table 8).

### Crystal structure of ^N828V^Pol2__CORE__ with a ribonucleotide in the active site

To explore why ^N828V^Pol ε incorporated more ribonucleotides than wild type Pol ε, we solved two different structures, namely ^N828V^Pol2_CORE_-dATP (PDB ID: 8b77) and ^N828V^Pol2_CORE_-UTP (PDB ID: 8b7E) at 2.70 Å and 2.60 Å resolution, respectively. The overall structure of ^N828V^Pol2_CORE_-dATP was identical to the wild-type Pol2_CORE_-dATP (PDB ID: 6qib) structure, and a comparison showed no significant structural changes in the active site (Supplementary Figure 4). The ^N828V^Pol2_CORE_-UTP structure showed a ribonucleotide bound to the active site in the presence of the steric gate, Y645, and showed that M644 provides stability to the active site in wild-type Pol ε. Compared to both the wild-type Pol2_CORE_-dATP and ^N828V^Pol2_CORE_-dATP structures, the presence of a ribonucleotide in the ^N828V^Pol2_CORE_ structure did not induce any significant structural changes to the residues surrounding the ribonucleotide (Figure 5a and Supplementary Figure 4). The 2’-OH group of the sugar was lifted slightly upwards and away from the Y645 plane, but without affecting the finger domain (Figure 5a) as V828 remained unchanged in its position. A comparison between ^N828V^Pol2_CORE_-UTP and ^M644G^Pol2_CORE_-UTP showed that the less bulky V828 could accept a ribonucleotide in the active site without introducing any strain in the finger domain or tilting of the base in the ribonucleotide (Figure 5b). In contrast, N828 shifted position and introduced a strain in the finger helix that might promote the opening of the finger domain followed by the dissociation of the ribonucleotide before a new bond is formed in the chemical step.

**Figure 5.**
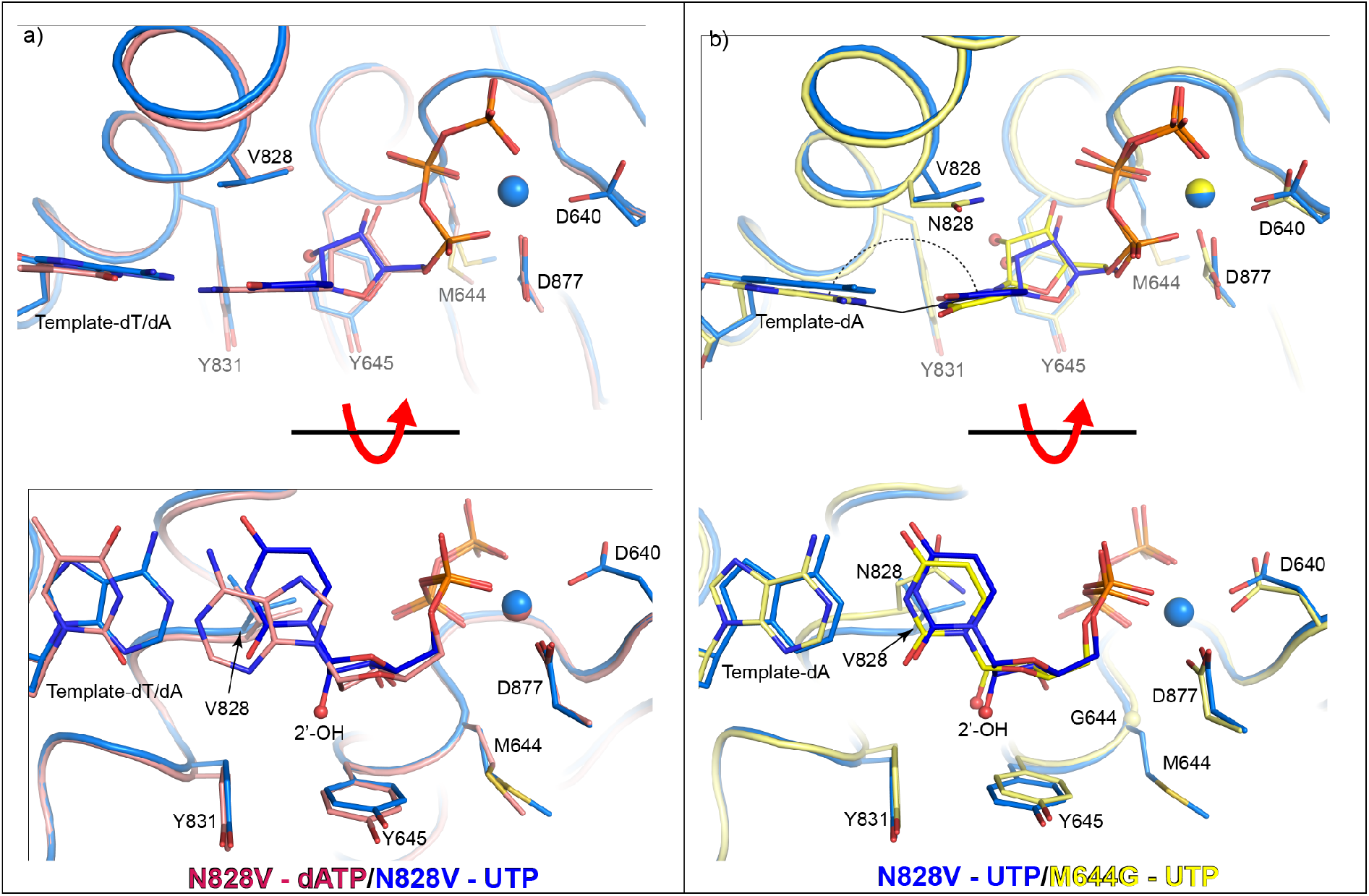
Crystal structures of ^N828V^Pol2_CORE_ with a ribonucleotide in the active site. a) The structures of ^N828V^Pol2_CORE_-UTP (in blue) and ^N828V^Pol2_CORE_-dATP (in pink) are superimposed. In contrast to the ^M644G^Pol2_CORE_-UTP structure, UTP is well aligned with the templating base, and the 2’-OH group (shown as a red sphere) has not moved sideways to avoid the clash with the tyrosine. Thus, the only structural difference between the UTP and the dATP is that the sugar moiety of UTP has been lifted upward compared to that of dATP. The view is rotated by 90° along the horizontal axis in the lower panel. b) The crystal structures of ^N828V^Pol2_CORE_ and ^M644G^Pol2_CORE_, both with UTP in the active site, are superimposed. Here, ^N828V^Pol2_CORE_-UTP is shown in blue and ^M644G^Pol2_CORE_-UTP is shown in yellow, and the position of the ribonucleotide differs between the two structures.

## DISCUSSION

Our data show that Pol ε discriminates NTPs through the concerted action of a steric gate residue (Y645) and a sensor in the finger domain (N828). Furthermore, these two mechanisms can independently of each other suppress ribonucleotide incorporation during DNA replication as demonstrated both *in vitro* and *in vivo* (Figures 3 and 4). Here, we will consider the generally accepted steric gate model, describing how B-family DNA polymerases discriminate NTPs based on their 2’-OH group, while presenting a refined model that also includes an asparagine that functions as a sensor and not, as suggested earlier, a polar filter (15) in the finger domain.

Biochemical studies of both high fidelity (6,12,51–53) and low fidelity (54,55) DNA polymerases have shown that substitution mutations for the steric gate residue lower sugar-based selectivity, with simultaneous reductions in the overall catalytic efficiency for incorporating dNTPs, thereby making it difficult to perform functional studies of steric gate mutants. For that reason, the neighboring amino acid, M644 in Pol ε, was substituted with the hope that it would destabilize the steric gate residue, Y645, in order to increase the incorporation of mis-matched nucleotides and the incorporation of ribonucleotides without severely affecting the catalytic efficiency (13,47). Fortunately, the ^M644G^Pol ε variant delivered a mutational signature that we were looking for and was also less efficient in discriminating against ribonucleotides (13,47). This ^M644G^Pol ε variant has since been widely used for studies that have explored the division of labor at the replication fork (14,16,56).

Here, the ^M644G^Pol ε variant allowed us to solve the first high-resolution structures of a B-family polymerase with a ribonucleotide in the polymerase active site. To our surprise, the position of the Y645 was unaltered. Instead, N828 in the finger domain, positioned on the opposite side of the nucleotide (not facing the 2’-OH), had moved 3.1 Å and a loose hydrogen bond with the β-phosphate was disrupted. The MD simulations of wild-type Pol ε corroborated that this was not an effect caused by the M644G substitution because N828 had the same favored positions in wild-type Pol ε when ATP was bound in the active site.

Based on our results, we present a refined model for how Pol ε discriminates ribonucleotides (Figure 6). A deoxyribonucleotide has a perfect fit and stacks on the steric gate residue while N828 in the finger domain forms a loose hydrogen bond with the β-phosphate of the deoxyribonucleotide. In contrast, the steric gate residue in the rigid palm domain induces an upward shift in the position of the sugar pucker of the bound ribonucleotide, and the altered nucleotide position does not allow N828 to form a non-essential loose hydrogen bond with the β-phosphate of the ribonucleotide while the unfavorable position of N828 induces a strain in the finger domain. As a result, phosphodiester bond formation is less likely to occur before the finger domain opens and the ribonucleotide dissociates. When replacing N828 with a valine, a ribonucleotide is accepted by the active site. Despite an upward shift in the position of the sugar moiety, there is no clash between the V828 and the 2’-C of the ribose that would induce a strain in the finger domain. Thus, the finger domain is likely to be maintained in a closed state increasing the probability that chemistry will occur with a ribonucleotide in the active site.

**Figure 6.**
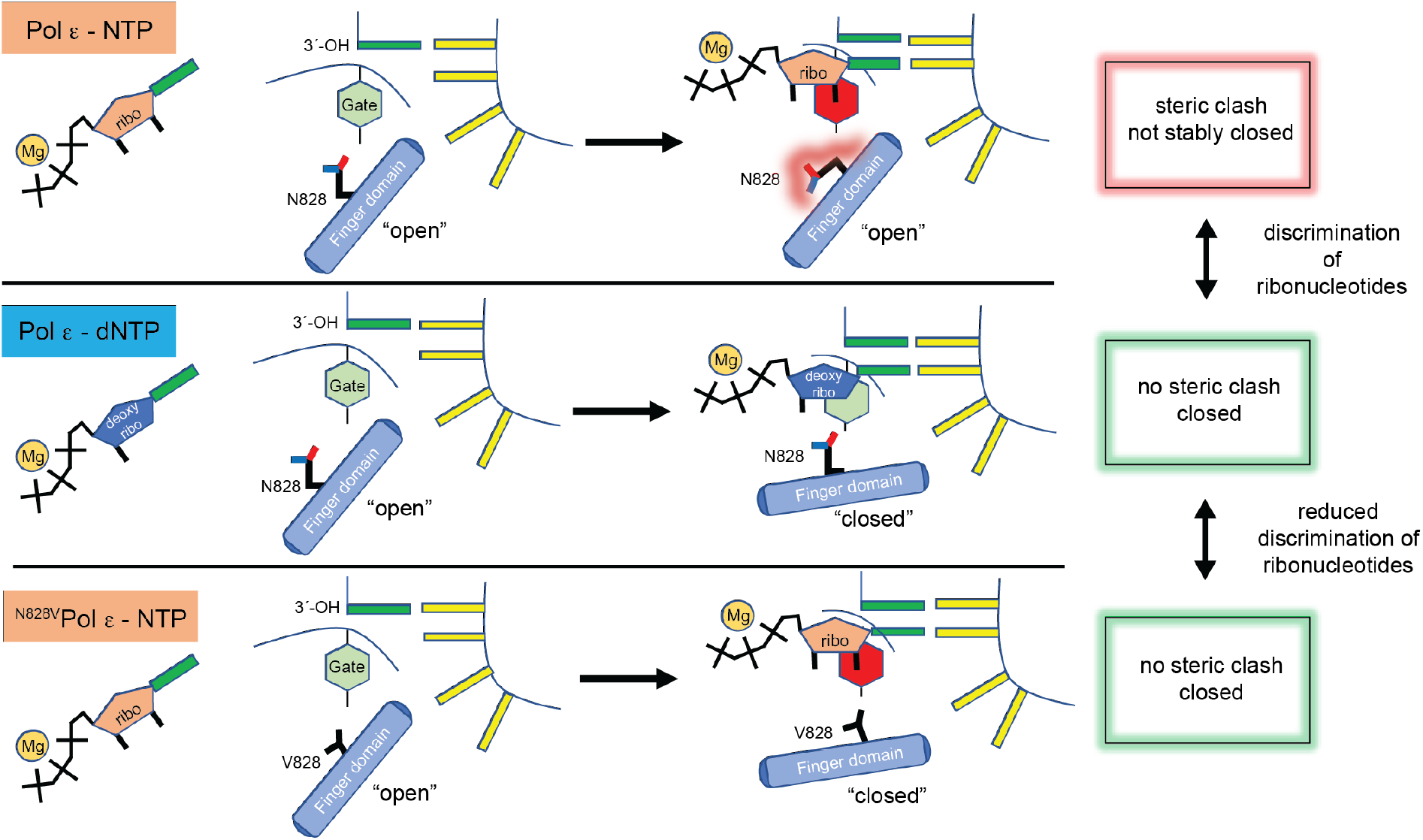
Model illustrating how the steric gate residue (tyrosine) and the sensor (asparagine) in the finger domain work together while discriminating NTPs from dNTPs. The red and green box refers to a steric clash between the sensor, N828, and the sugar moiety of the bound nucleotide. In the bottom panel, the N828V substitution allows the finger to close as there is no steric clash with the sugar moiety, despite the clash between the 2’-OH and the steric gate residue.

We propose that the sensor recognize a ribonucleotide by steric hindrance and this model differ from the earlier described polar filter that was proposed to pull the nucleotide closer to the enzyme surface *via* hydrogen bonds to the 3’-OH group and triphosphate of the incoming nucleotide (15). To compare the two models, we revisited the structures that were described in the original article, focusing on the hydrogen bonding interactions upon which the “polar filter” model depends (15). A combination of both visual inspection of the proposed hydrogen bonds, and analysis of hydrogen bonding interactions in each structure using ChimeraX’s H-bond tool 1. (57) (technical details of how H-bonds are defined are described in Supplementary Table 9) recognizes only a few (3/20) of the proposed H-bonds as strict hydrogen bonds, a further 9/20 interactions are only identified as hydrogen bonds if the definition of a hydrogen bond is relaxed, and the remainder (8/20) do not fulfill even relaxed hydrogen bond criteria (in one case, the interaction is indirect, *via* an intervening water molecule). In general, the hydrogen bonds were not recognized due to a too long distance and/or an impossible angle between donor and acceptor for the hydrogen bond. This was true in structures of both Y-family polymerases and B-family polymerases on which the polar filter model was based. Therefore, in the absence of strong or feasible hydrogen bonds, we argue that steric hindrance affects the closing of the finger and this is a more conceivable mechanism for suppression of ribonucleotide incorporation.

Both our *in vitro* and *in vivo* results support our proposed sensor model because ^N828V^Pol ε more frequently incorporated ribonucleotides at physiological concentrations of dNTPs and NTPs. Thus, ^N828V^Pol ε might be a useful alternative to ^M644G^Pol ε for addressing questions related to the division of labor at the replication fork. Interestingly, based on our results M644G and N828V differ both with sequence contexts where ribonucleotides are more likely to be incorporated and the frequency at which specific mis-incorporations are made. Thus, depending on the question to be addressed, M644G and N828V might serve as controls for each other and might also be more or less suitable for different experiments.

We then asked whether combining the M644G and N828V substitutions could convert Pol ε into an RNA polymerase when only given ribonucleotides. The ^M644G-N828V^Pol ε variant was capable of synthesizing a short stretch of RNA (4 nucleotides in 80 seconds) in the presence of proofreading activity and under single-hit conditions. However, when given physiological concentrations of dNTPs and NTPs, dNTPs were preferred by the active site although at least 1 ribonucleotide was incorporated per 30 nucleotides (compared with wild-type Pol ε at 1 per 1250 nucleotides(58) or ^M644G^Pol ε at 1 per 91 nucleotides (13)). Thus, the ^M644G-N828V^Pol ε variant incorporates unprecedented levels of ribonucleotides *in vivo*, likely saturating the ribonucleotide excision repair system (59). The high load of ribonucleotides in the genome is the likely mechanism behind the strong growth inhibition of haploid strains expressing ^M644G-N828V^Pol ε (Supplementary Figure 3a). This growth defect disqualifies ^M644G-N828V^Pol ε from further *in vivo* studies.

A comparison with other B-family DNA polymerases showed that both Y645 and N828 are structurally conserved residues (Supplementary Figure 5), thus reinforcing their functional importance. Thus, we propose that the mechanism by which Pol ε discriminates against ribonucleotides might be conserved among all B-family polymerases. Interestingly, Beese and co-workers found that the *Bacillus* A-family DNA polymerase I also depends on the closing of the finger domain for discriminating ribonucleotides (5). In fact, the structure of the A-family polymerase could only be obtained with a ribonucleotide in the active site after substituting a tyrosine for a phenylalanine at the same position where N828 is located in the finger domain in Pol ε. Thus, other family A, X, and Y polymerases should be revisited to explore if they have a built-in sensor that comes into play when there is a clash between the 2’-OH and a steric gate residue.

## DATA AVAILABILITY

Structures are deposited in the Protein Data Bank under accession codes PDB ID 8b76 (M644G-dTTP), 8b6k (M644G-dCTP), 8b79 (M644G-UTP), 8b67 (M644G-CTP), 8b77 (N828V-dATP), and 8b7e (N828V-UTP). All simulation starting structures, representative input files, non-standard parameter files, and snapshots from our simulations are deposited at Zenodo (DOI: 10.5281/zenodo.7446658)

## SUPPLEMENTARY DATA STATEMENT

Supplementary Data are available at NAR online.

## AUTHOR CONTRIBUTIONS

V.P., G.O.B., and P.O. purified the proteins; G.O.B. and P.O. carried out the biochemical experiments; V.P. crystallized the proteins and solved and refined all structures; Y.K. performed the MD simulations; G.O.B. performed all genetics experiments; and V.P., Y.K, S.C.L.K., and E.J. conceived the research and wrote the article. All authors discussed the results and commented on the manuscript.

## AUTHOR INFORMATION

The authors declare no competing financial interests. Correspondence and requests for materials should be addressed to E.J.

## ACKNOWLEDGEMENTS

We thank Alexandre Barrozo for help with establishing the system for Pol2 in the molecular dynamics simulations. Experiments were performed on beamlines ID29 and ID30A-3 at the European Synchrotron Radiation facility (ESRF), Grenoble. We are grateful to the local contact at ESRF for providing assistance in using beamlines ID29 and ID30A-3. We also acknowledge the MAX IV Laboratory for time on the BioMAX beamline under Proposal 20180236. Research conducted at MAX IV, a Swedish national user facility, is supported by the Swedish Research council under contract 2018-07152, the Swedish Governmental Agency for Innovation Systems under contract 2018-04969, and Formas under contract 2019-02496. Finally, we acknowledge DESY (Hamburg, Germany), a member of the Helmholtz Association HGF, for the provision of experimental facilities. Parts of this research were carried out at P13-PETRA III, and we would like to thank Drs Johanna Hakanpää and Saravanan Panneerselvam for assistance in using the P13 beamline. Beamtime was allocated for proposal 2019-MX-696.

## FUNDING

This work was supported by the Knut and Alice Wallenberg Foundation [grant numbers 2018.0140, 2019.0431] (S.C.L.K), the Swedish Research Council [grant number 2019-03499] (S.C.L.K), [grant number 2021-01104] (E.J) and the Swedish Cancer Society (E.J.). The computations were enabled by resources provided by the Swedish National Infrastructure for Computing (SNIC), which is partially funded by the Swedish Research Council through grant agreement no. 2018-05973.

## CONFLICT OF INTEREST

The authors declare no conflicts of interest.

## Notes

### Competing Interest Statement

The authors have declared no competing interest.

