## Supplementary material for "A sensor complements the steric gate when DNA polymerase ε discriminates ribonucleotides"

### Table of Contents

**Supplementary Figures 1-7…………..……………………………………………………………S3-S8**

Supplementary Figure 1. Comparison of the polymerase active site in wild-type $Pol2_{CORE}$ and $^{M644G}Pol2_{CORE}$ ternary complexes with a dNTP. ..........................................................................................................S3

Supplementary Figure 2. Structural comparison of $^{M644G}Pol2_{CORE}$-CTP and $^{M644G}Pol2_{CORE}$-dCTP………………………………………………..………………………………………...S4

Supplementary Figure 3. A tetrad analysis shows that haploids with a *pol2-M644G,N828V* allele has a strong growth defect and that *rnh201Δ, pol2-N828V* strains are viable…………………………………………S5

Supplementary Figure 4. Crystal structure of $^{N828V}Pol2_{CORE}$ with an incoming dATP, superimposed on the structure of wild-type $Pol2_{CORE}$-dATP……………………………………....……………………S6

Supplementary Figure 5. N828 in Pol ε is a structurally conserved asparagine in B-family DNA polymerases.S7

Supplementary Figure 6. Box plots showing distribution of key distances and angles measured in MD simulations………………………………………………………………………………………………S8

Supplementary Figure 7. Box plots showing distribution of metal-ligand distances (Å) during our simulations………………………………………………………………………………………………S8

**Supplementary Tables 1-9……………………………..………………………………………S9-S13**

Supplementary Table 1. Data collection and refinement statistics ……….....………………………………..S9

Supplementary Table 2. Distance (Å) between $C_\alpha$-atom of D877 and the OH group oxygen of Y645 during simulations of wild-type and M644G Pol ε in complex with dATP and ATP………….……...…………..…S10

Supplementary Table 3. Distance (Å) between the 2'-C atom of the bound nucleotide (dATP/ATP) and the $C_{\delta 1}$ atom of Y645 during simulations of wild-type and M644G Pol ε in complex with dATP and ATP……..…………………………………………………………………....……………………………..S10

Supplementary Table 4. Angle (°) between the O2B atom of the dNTP and the $C_\alpha$ and $C_\gamma$ atoms of the N828 side chain during simulations of wild-type and M644G Pol ε in complex with dATP and ATP…………….S10

Supplementary Table 5. Force field parameters for the tetrahedral $Zn^{2+}$ dummy model used in this work ……….....…………………………………………………………………………………………………S11

Supplementary Table 6. Metal-ligand distances (Å) between Zn metal center and its coordinating cysteine sidechain sulfur atoms calculated from out molecular dynamics simulations of wild-type and M644G Pol ε in complex with dATP and ATP……………………………………………………………………………S11

Supplementary Table 7. Spontaneous mutation rates …………………....…………..………………..…S12

Supplementary Table 8. Mutations found in *CAN1* gene of $Can^R$ yeast strains……………………………...S12

Supplementary Table 9. Analysis of hydrogen bonds on which the "polar filter" model depends on………..S13

**Supplementary References…………………………..………………………………………….S14**

#### Supplementary Figures

Supplementary figure 1

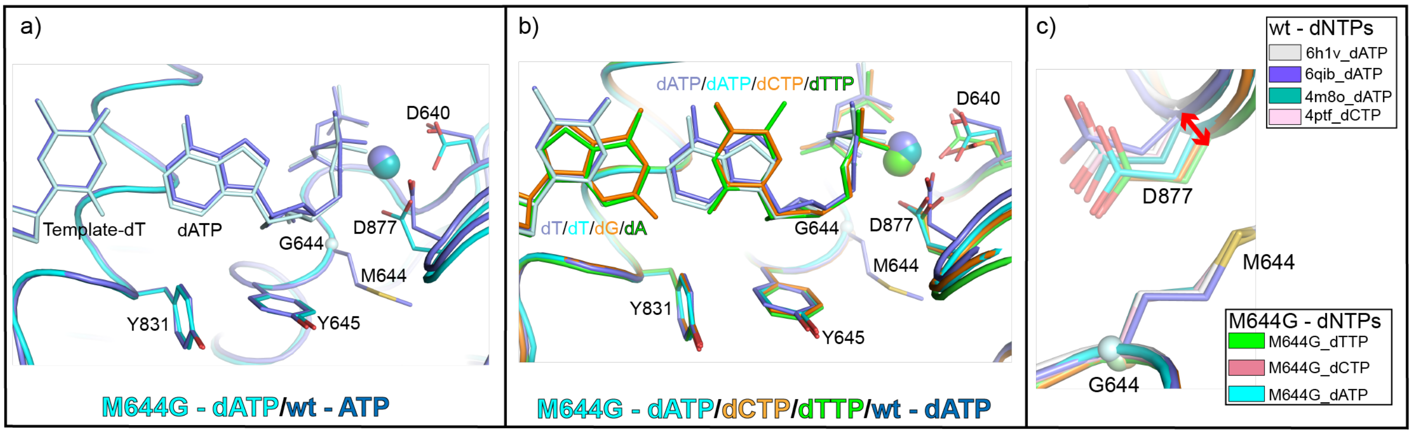

**Supplementary Figure 1. Comparison of the polymerase active site in wild-type Pol2$_{CORE}$ and $^{M644G}$Pol2$_{CORE}$ ternary complexes with a dNTP.** a) The structures of $^{M644G}$Pol2$_{CORE}$ (cyan) and wild-type Pol2$_{CORE}$ (blue) with dATP bound in the active site are superimposed. b) All $^{M644G}$Pol2$_{CORE}$ structures containing dATP (cyan), dCTP (orange), or dTTP (green) are superimposed with the wild-type Pol2$_{CORE}$ (blue) structure. c) The structures of $^{M644G}$Pol2$_{CORE}$ with different dNTPs (dATP, dCTP, or dTTP) are aligned to the available wild-type Pol2$_{CORE}$ structures (PDB IDs: 6h1v (1), 6qib (1), 4m8o (2), and 4ptf (3)). The loop containing D877 is slightly shifted (~ 1Å) in all $^{M644G}$Pol2$_{CORE}$-dNTP structures when compared to the wild-type Pol2$_{CORE}$-dNTP structures.

Supplementary figure 2

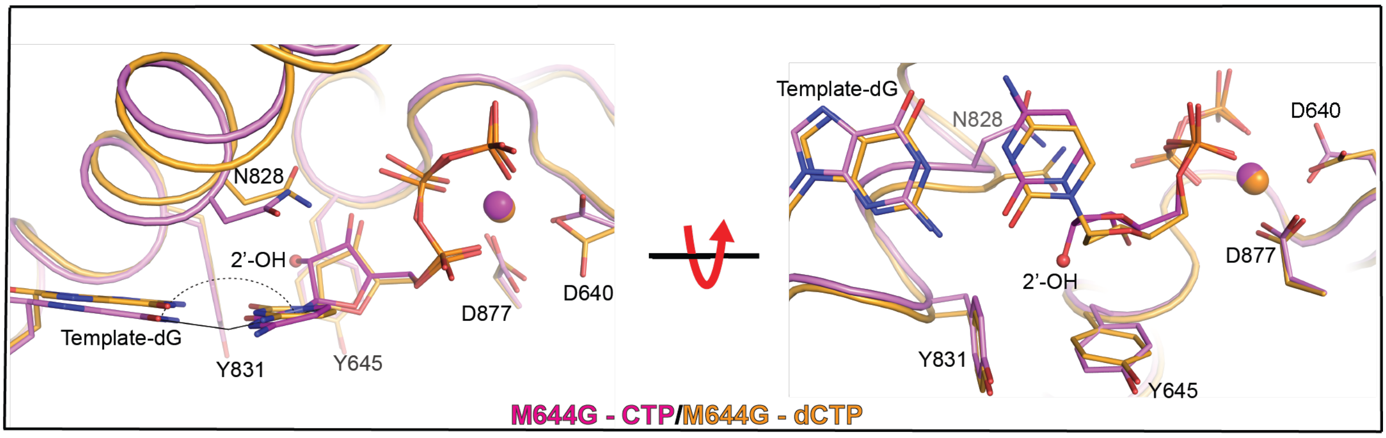

**Supplementary Figure 2. Structural comparison of $^{M644G}$Pol2$_{CORE}$-CTP and $^{M644G}$Pol2$_{CORE}$-dCTP.** a) Comparison of $^{M644G}$Pol2$_{CORE}$-CTP (purple) and $^{M644G}$Pol2$_{CORE}$-dCTP (orange) showing that the ribonucleotide takes a slightly different position compared to the deoxyribonucleotide. The nonplanar alignment with the templating base (dG) and CTP and the hydrogen bond between N828 and the beta-phosphate of dTTP are highlighted. b) The view is rotated by 90° along the horizontal axis to show the position of the 2'-OH group relative to the steric gate, Y645, and the shift of N828 in the finger domain.

#### Supplementary figure 3

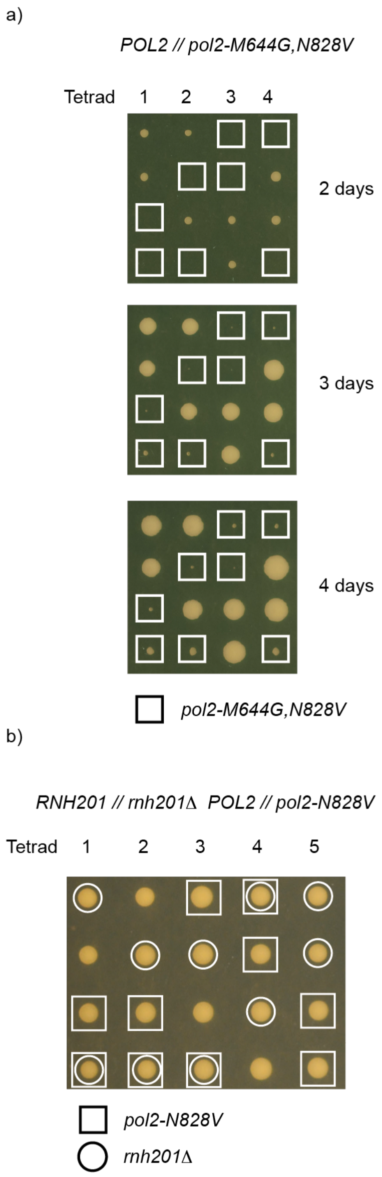

**Supplementary Figure 3. Tetrad analysis showing that haploids with a *pol2-M644G,N828V* allele have a strong growth defect and that *rnh201Δ, pol2-N828V* strains are viable.** a) *POL2/pol2-M644G,N828V* diploid yeast in the E134 strain background were sporulated followed by tetrad dissection. Colonies with cells carrying the *pol2-M644G,N828V* allele are in boxes. b) Diploid yeast, *RNH201/rnh201Δ, POL2/pol2-N828V* in the E134 strain background were sporulated followed by tetrad dissection.

Supplementary figure 4

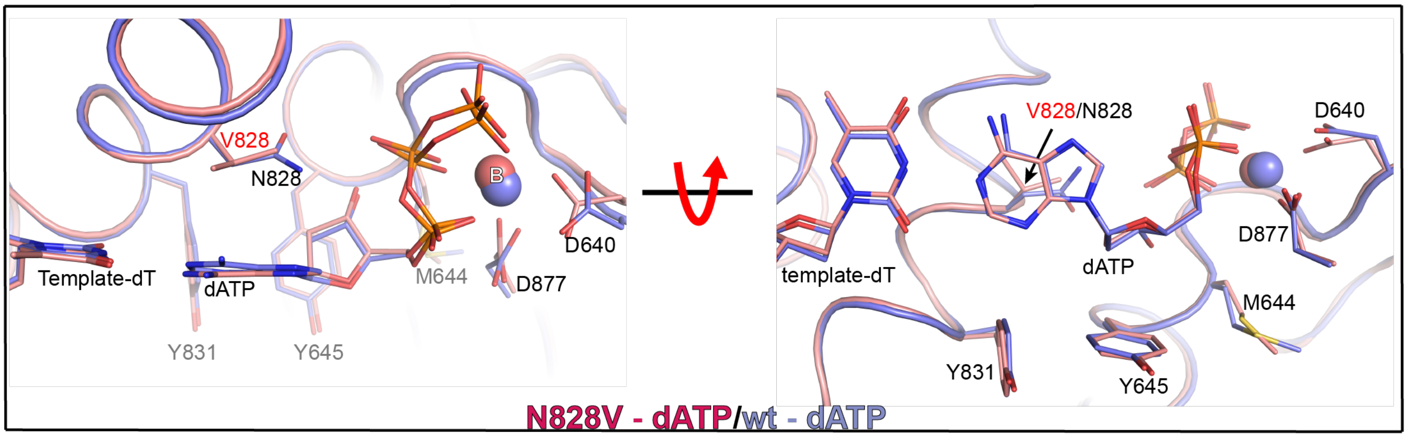

**Supplementary Figure 4. Crystal structure of $^{N828V}$Pol2$_{CORE}$ with an incoming dATP superimposed on the structure of wild-type Pol2$_{CORE}$-dATP.** a) Comparison of $^{N828V}$Pol2$_{CORE}$-dATP (in pink) and wild-type Pol2$_{CORE}$-dATP (in blue) showing that the deoxyribonucleotide had an identical position despite the N828V substitution in the finger domain. b) The view is rotated by 90° along the horizontal axis.

Supplementary figure 5

a)

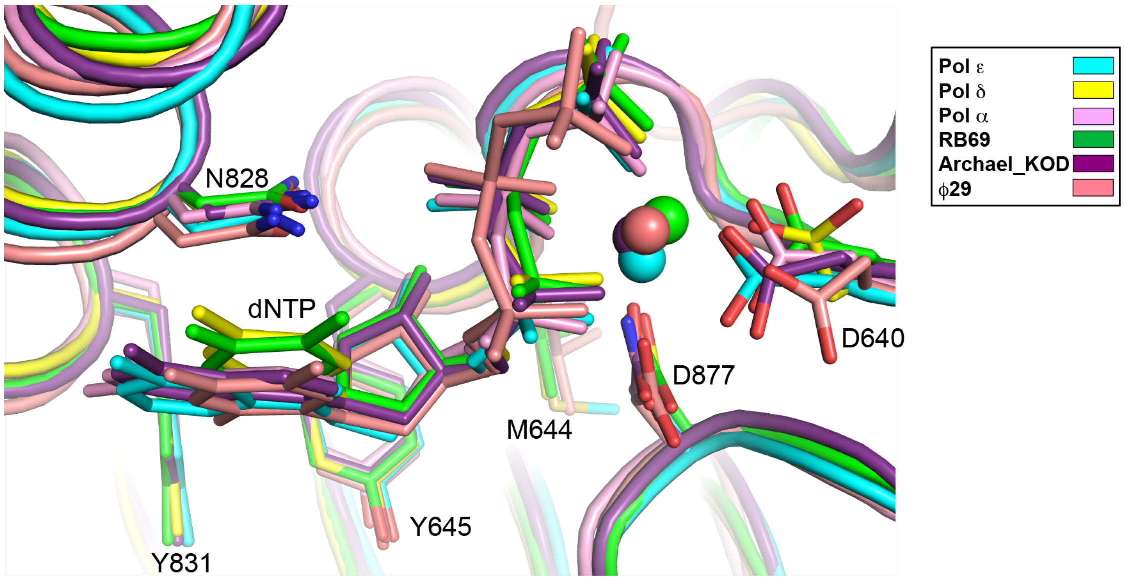

b)

| Pol ε residue | 640 ... 644 645 | 828 ... 831 | 877 |
|---|---|---|---|
| Pol ε | Y H V D V A S M Y P N I M T | Q L A H K V I L N S F Y G Y | E L D T D G I W |
| Pol δ | A T L D F N S L Y P S I M M | Q L A L K I S A N S V Y G F | Y G D T D S V M |
| Pol α | L V M D F N S L Y P S I I Q | Q Q A L K L T A N S M Y G C | Y G D T D S V M |
| RB69 | M S F D L T S L Y P S I I R | Q I N R K L L I N S L Y G A | Y G D T D S I Y |
| Archeal KOD | V Y L D F R S L Y P S I I I | Q R A I K I L A N S Y Y G Y | Y S D T D G F F |
| Pfu | V Y L D Y K S L Y P S I I I | Q K A I K L L A N S F Y G Y | Y I D T D G L Y |
| P. abyssi | V S L D F R S L Y P S I I I | Q R A I K I L A N S Y Y G Y | Y I D T D G L Y |
| SsPolB1 | T V L D F A S L Y P S I I R | Q R A M K V F I N A T Y G V | Y G D T D S L F |
| ϕ29 | M V F D V N S L Y P A Q M Y | Q L A - K L M L N S L Y G K | Y C D T D S I H |

**Supplementary Figure 5. N828 in Pol ε is a structurally conserved asparagine in B-family DNA polymerases.** a) Superposition of structures from B-family DNA polymerases with a deoxyribonucleotide in the active site and a metal coordinated by catalytic aspartates in the B-site. b) Structure-based sequence alignment of B-family DNA polymerases highlighting the residues shown in the structural superposition (see panel a).

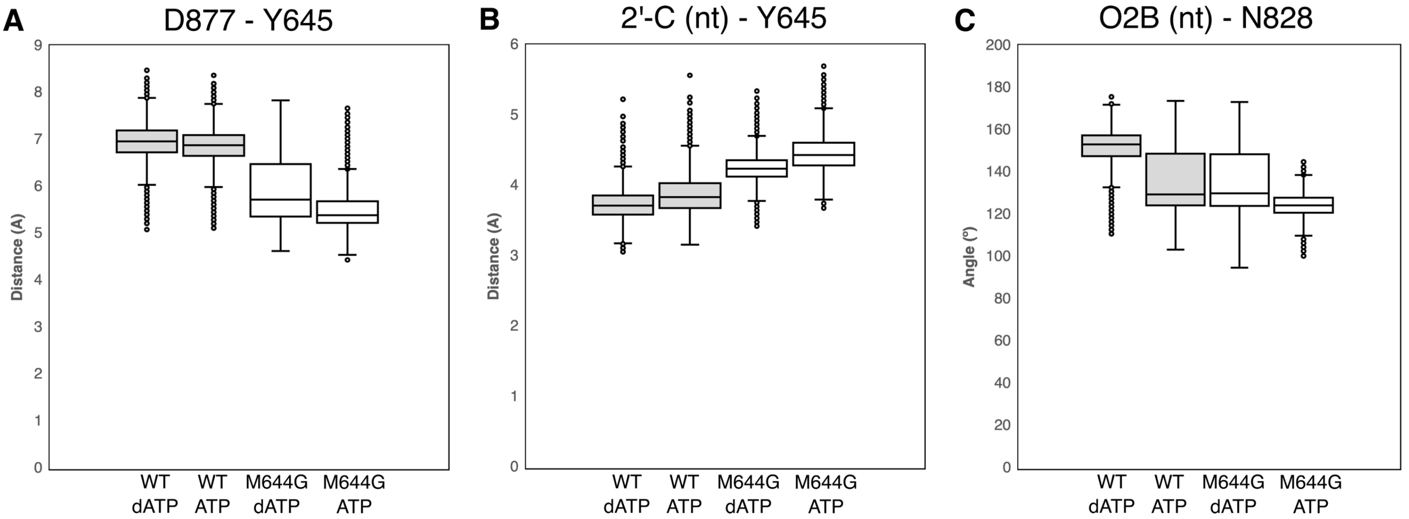

**Supplementary Figure 6**. **Box plots showing distribution of key distances and angles measured in MD simulations.** a) Distance between the $C_\alpha$-atom of D877 and the OH-group oxygen of Y645. b) Distance between the 2'-C atom of the bound nucleotide (dATP/ATP) and the $C_{\delta 1}$ atom of Y645. c) Angle between the O2B atom of the nucleotide, and the $C_\alpha$ and $C_\gamma$ atoms of the N828 side chain. All distances are in Å and angles in degrees (°). Shaded and white boxes correspond to wild-type Pol ε and M644G Pol ε respectively. Whiskers indicate standard deviations. Average and standard deviation values corresponding to panels a - c are provided in Supplementary Tables 2 through 4 respectively.

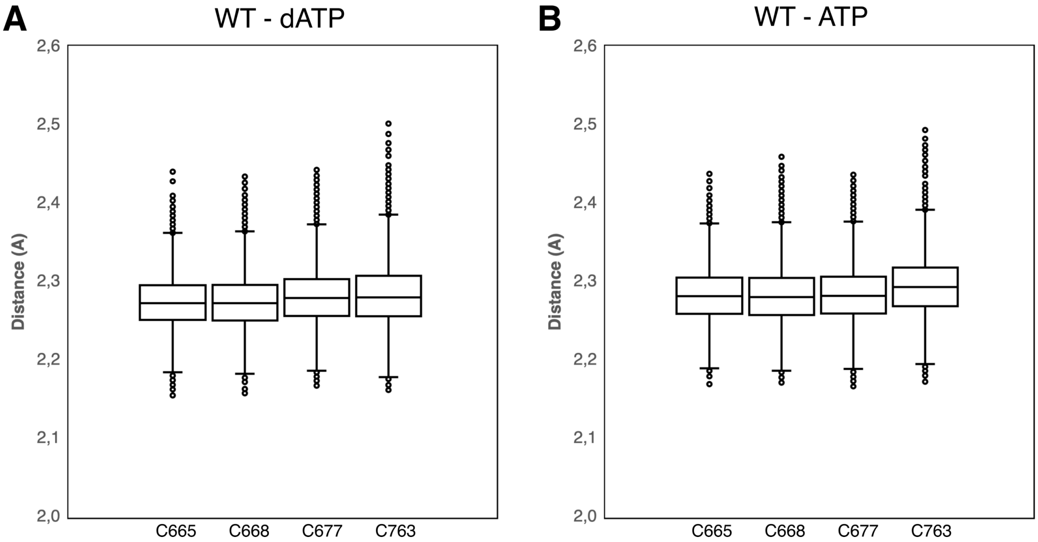

**Supplementary Figure 7**. **Box plots showing distribution of metal-ligand distances (Å) during our simulations.** Shown here are distances between the Zn metal center and its coordinating cysteine sidechain sulfur atoms, calculated from MD simulations of wild-type Pol ε in complex with a) dATP and b) ATP**.** Average and standard deviation values corresponding to panels a and b are provided in Supplementary Table 6.

#### Supplementary Tables

Supplementary Table 1. Data collection and refinement statistics

| **PDB ID code** | 8b76 | 8b6k | 8b79 | 8b67 | 8b77 | 8b7e |
|---|---|---|---|---|---|---|
| Structure | M644G-dTTP | M644G-dCTP | M644G-UTP | M644G-CTP | N828V-dATP | N828V-UTP |
| **Data Collection** | | | | | | |
| Spacegroup | P2 | P2 | P2 | C2 | C2 | P2 |
| Cell dimensions | | | | | | |
| a(Å) | 154.0 | 154.4 | 154.4 | 162.1 | 159.0 | 154.4 |
| b(Å) | 69.8 | 70.4 | 70.7 | 67.4 | 71.03 | 70.0 |
| c(Å) | 159.0 | 158.6 | 159.4 | 152.1 | 155.25 | 158.3 |
| α, β, γ (°) | 90.0, 112.6, 90.0 | 90.0, 112.7, 90.0 | 90.0, 112.8, 90.0 | 90.0, 111.5, 90.0 | 90.0, 113.0, 90.0 | 90.0, 112.6, 90.0 |
| Resolution (Å) | 2.60 | 2.50 | 2.65 | 2.60 | 2.70 | 2.60 |
| R-merge (%) | 9.2 (68.4) | 20 (126) | 12.7 (172.6) | 14.9 (49) | 12.3 (69.5) | 16.6 (124.5) |
| I/sigmaI | 7.4 (1.5) | 5.4 (1.1) | 9.9 (1.0) | 5.6 (2.7) | 4.5 (2.0) | 6.8 (1.4) |
| Completeness | 99.5 (99.8) | 99.8 (99.1) | 99.8 (99.9) | 98.5 (99.8) | 98.6 (99.7) | 98.9 (99.9) |
| CC(1/2) | 0.99 (0.42) | 0.98 (0.08) | 0.99 (0.46) | 0.96 (0.40) | 0.99 (0.42) | 0.99 (0.32) |
| Redundancy | 3.0 (3.1) | 3.9 (3.6) | 6.4 (6.6) | 3.7 (3.9) | 4.2 (4.4) | 4.3 (4.4) |
| **Refinement** | | | | | | |
| Resolution (Å) | 2.6 | 2.5 | 2.65 | 2.6 | 2.7 | 2.6 |
| Reflections | 96114 | 109009 | 92507 | 46654 | 43315 | 95325 |
| $R_{work}/R_{free}$ (%) | 22.0/24.6 | 22.9/26.0 | 21.7/24.8 | 23.9/27.1 | 23.9/27.2 | 20.0/23.0 |
| TLS groups | - | 7 | - | 11 | 6 | 19 |
| Atoms | | | | | | |
| Macromolecules | 18761 | 18642 | 19007 | 9097 | 9293 | 18674 |
| Ligands | 72 | 58 | 72 | 36 | 31 | 68 |
| $H_2O$ | 53 | 0 | 12 | 15 | 0 | 82 |
| Average B-factors | | | | | | |
| Macromolecules | 64.4 | 59.9 | 76.6 | 56.3 | 83.3 | 58.2 |
| Ligands | 48.0 | 34.6 | 59.5 | 37.6 | 45.2 | 41.2 |
| $H_2O$ | 47.1 | - | 65.6 | 36.7 | - | 43.6 |
| RMSD | | | | | | |
| Bond length (Å) | 0.004 | 0.004 | 0.005 | 0.005 | 0.005 | 0.006 |
| Bond angles (°) | 0.69 | 0.66 | 1.003 | 0.88 | 0.86 | 0.88 |
| Ramachandran | | | | | | |
| Favoured (%) | 96.2 | 96.1 | 96.0 | 96.7 | 94.7 | 96.3 |
| Allowed (%) | 3.7 | 3.9 | 4.0 | 3.3 | 5.3 | 3.6 |

Supplementary Table 2. Distance (Å) between the $C_\alpha$-atom of D877 and the OH-group oxygen of Y645 during MD simulations of wild-type and M644G Pol ε in complex with dATP and ATP.[a]

| WT | | M644G | |
|---|---|---|---|
| **dATP** | **ATP** | **dATP** | **ATP** |
| 7.0 ± 0.4 | 6.9 ± 0.4 | 5.9 ± 0.6 | 5.5 ± 0.4 |

[a] All distances are average values and standard deviations, averaged over 3 production replicas of 100 ns simulation time each, as described in the Materials and Methods, with snapshots taken every 12.5 ps for the analysis.

Supplementary Table 3. Distance (Å) between the 2'-C atom of the bound nucleotide (dATP/ATP) and the $C_{\delta 1}$ atom of Y645 during MD simulations of wild-type and M644G Pol ε in complex with dATP and ATP.[a]

| WT | | M644G | |
|---|---|---|---|
| **dATP** | **ATP** | **dATP** | **ATP** |
| 3.7 ± 0.2 | 4.3 ± 0.2 | 3.9 ± 0.3 | 4.5 ± 0.2 |

[a] All distances are average values and standard deviations, averaged over 3 production replicas of 100 ns simulation time each, as described in the Materials and Methods, with snapshots taken every 12.5 ps for the analysis.

Supplementary Table 4. Angle (°) between the O2B atom of the dNTP and the $C_\alpha$ and $C_\gamma$ atoms of the N828 side chain during simulations of wild-type and M644G Pol ε in complex with dATP and ATP.[a]

| WT | | M644G | |
|---|---|---|---|
| **dATP** | **ATP** | **dATP** | **ATP** |
| 150.9 ± 10.5 | 135.0 ± 13.7 | 134.8 ± 13.8 | 124.5 ± 5.4 |

[a] All distances are average values and standard deviations, averaged over 3 production replicas of 100 ns simulation time each, as described in the Materials and Methods, with snapshots taken every 12.5 ps for the analysis.

Supplementary Table 5. Force field parameters for the tetrahedral $Zn^{2+}$ dummy model used in this work.[a]

| **Bonded Interactions** | | | | |
|---|---|---|---|---|
| **Bond type[a]** | | | $\boldsymbol{K_b}$ | $\boldsymbol{r_0}$ |
| Zn-D | | | 800.0 | 0.900 |
| **Angle type[b]** | | | $\boldsymbol{K_\theta}$ | $\boldsymbol{\theta_0}$ |
| D-Zn-D | | | 250.0 | 109.5 |
| **Mass (*m*), Charge (*e*), and Non-bonded Interactions[c]** | | | | |
| **Atom Type** | $\boldsymbol{m}$ | $\boldsymbol{e}$ | $\boldsymbol{A_i}$ | $\boldsymbol{B_i}$ |
| Zn | 53.39 | –1.00 | 68.00 | 38.00 |
| D | 3.00 | 0.75 | 0.05 | 0.00 |

[a] Zn represents the zinc metal center atom, while D represents the dummy atoms. [a] $U_b = K_b(b - b_0)^2$, where $K_b$ is in kcal mol$^{-1}$ Å$^{-2}$ and $r_0$ is in Å. [b] $U_\theta = (1/2)K_\theta(\theta - \theta_0)^2$, where $K_\theta$ is in kcal mol$^{-1}$ rad$^{-2}$ and $\theta_0$ is in degrees. [c] Lennard–Jones parameters are given in units of [kcal$^{1/2}$ mol$^{-1/2}$ Å$^6$] for $A_i$ and [kcal$^{1/2}$ mol$^{-1/2}$ Å$^3$] for $B_i$.

Supplementary Table 6. Metal-ligand distances (Å) between the Zn metal center and its coordinating cysteine sidechain sulfur atoms calculated from our MD simulations of wild-type Pol ε in complex with dATP and ATP. Experimental distances for the corresponding atom pairs have been measured from the crystal structure coordinates of wild-type Pol ε (PDB: 4m8o).[a]

| **Residue$^{\text{atom}}$** | **Crystal Structure** | **dATP[a]** | **ATP[a]** |
|---|---|---|---|
| C665$^{SG}$ | 2.3 | 2.3 ± 0.04 | 2.3 ± 0.04 |
| C668$^{SG}$ | 2.3 | 2.3 ± 0.04 | 2.3 ± 0.04 |
| C677$^{SG}$ | 2.2 | 2.3 ± 0.04 | 2.3 ± 0.04 |
| C763$^{SG}$ | 2.4 | 2.3 ± 0.04 | 2.3 ± 0.04 |

[a] All distances are average values and standard deviations, averaged over 3 production replicas of 100 ns simulation time each, as described in the Materials and Methods, with snapshots taken every 12.5 ps for the analysis.

Supplementary Table 7. Spontaneous mutation rates.[a]

| Genotype | $CAN^r$ | |
|---|---|---|
| | **Mutation rate (x$10^{-7}$)** | **Relative [a)]** |
| Wild-type | 4.6 (4.0-5.9) | 1 |
| *pol2-N828V* | 17.5 (14.4-20.1) | 3.8 |
| *msh6Δ* | 16.6 (14.8- 25.5) | 3.6 |
| *pol2-N828V msh6Δ* | 642 (553-916) | 140 |

[a] Rates are relative to the wild-type strain. Measurements were performed as described in the Methods. Values in parentheses are 95% confidence intervals, calculated as described in ref. (4).

Supplementary Table 8. Mutations found in the *CAN1* gene of $Can^R$ yeast strains.

| **Types of mutations** | **Number of mutations in the different strains** | | | | **Rates (×$10^{-8}$)** | | | |
|---|---|---|---|---|---|---|---|---|
| | **Wild type** | ***Pol2-N828V*** | ***Msh6Δ*** | ***Pol2-N828V Msh6Δ*** | **Wild type** | ***Pol2-N828V*** | ***Msh6Δ*** | ***Pol2-N828V Msh6Δ*** |
| Base substitutions: | | | | | | | | |
| AT->TA | 1[a] | 74[b] | 1 | 4 | 1 | 139 | 2 | 265 |
| AT->CG | 2 | 3 | 6 | 1 | 2 | 6 | 11 | 66 |
| AT->GC | 3 | 2 | 6 | 1 | 3 | 4 | 11 | 66 |
| GC->CG | 12 | 0 | 3 | 2 | 12 | <1 | 5 | 132 |
| GC->AT | 7 | 5 | 52 | 55 | 7 | 9 | 92 | 3642 |
| GC->TA | 10 | 3 | 18 | 32 | 10 | 6 | 32 | 2119 |
| –1 deletion | 5 | 2 | 5 | 1 | 5 | 4 | 9 | 66 |
| +1 insertion | 2 | 2 | 1 | 1 | 2 | 4 | 2 | 66 |
| Complex mutation | 1 | | | | 1 | <1 | <1 | <1 |
| Other | 5 | 2 | 2 | 0 | 5 | 4 | 3 | <1 |
| Total | 48 | 93 | 94 | 97 | | | | |

[a] Data from Aksenova *et al*. (5). [b] 63 out of 74 substitutions were A->T, thus leading to an imbalance that could result from T-T mismatches during leading strand synthesis.

Supplementary Table 9. Exploring the existence of the possible hydrogen bonds on which the “polar filter” model depends (6). The analysis was performed using the H-bonds tool in ChimeraX(7).

| Y-family Polymerase | PDB ID | Finger residue – O3’ hydrogen bond[a] | Finger residue – beta phosphate hydrogen bond[a] |
|---|---|---|---|
| PolK (*H. sapiens*) | 2oh2 | Loose | None |
| Rev1 (*S. cerevisiae*) | 2aq4 | Loose | Strict |
| PolI (*H. sapiens*) | 2dpj | None | Strict |
| Dpo4 (*S. solfataricus*) | 4qw8 | None | Strict |
| B-family polymerases | | | |
| Phi29 DNA Polymerase (phage) | 2pyl | none (indirect via water molecule) | none (indirect via water molecule) |
| Pol δ (*S. cerevisiae*) | 3iay | Loose | Loose |
| PolII (*E. coli*) | 3k57 | Loose | Loose |
| RB69 DNA polymerase (phage) | 4fjl | Loose | Loose |
| Pol ε (*S. cerevisiae*) | 4m8o | None | Loose |
| Pol α (*H. sapiens*) | 4qcl | None | None |

a) Here, we separated “strict” hydrogen bonds (fulfilling the precise geometric criteria for a hydrogen bond) from “loose” hydrogen bonds, identified with relaxed hydrogen-bond constraints (by 0.4 Ångströms and 20.0 degrees) indicating that tolerances have been applied to the precise geometric criteria. These standard criteria in ChimeraX came from a survey of high-resolution structures(8); empirically, tolerances of 0.4 Ångström and 20.0 degrees are considered to work well for most macromolecular structures. “None” indicates that ChimeraX could not find a hydrogen bond even with the relaxed hydrogen-bond restraints.
